# BertMS-enabled molecular networking for unknown compounds dereplication

**DOI:** 10.64898/2026.03.19.712370

**Authors:** Luning Zhou, Shuang Wu, Jixing Peng, Xiaofei Huang, Wenxue Wang, Dehai Li

## Abstract

Spectral similarity is widely used as a proxy for structural similarity in tandem mass spectrometry (MS/MS) analyses, including library matching and molecular networking. However, the relationship between spectral similarity scores and true structural similarity remains imperfect, limiting compound identification in metabolomics studies. Here, we present BertMS, a spectral similarity framework based on bidirectional encoder representations from transformers (BERT), which learns contextualized representations of fragment ions from large-scale MS/MS data. Using datasets from MoNA and GNPS comprising over 100,000 unique molecules, we systematically evaluate BertMS against existing methods, including cosine similarity and Spec2Vec. BertMS shows improved performance across multiple evaluation metrics, with average gains of approximately 15–25% depending on the task. Notably, improvements are most evident in molecular similarity assessment. We further demonstrate the applicability of BertMS in molecular networking and dereplication of microbial metabolites, where it enables more consistent identification of structurally related compounds. Together, these results demonstrate that transformer-based representations improve spectral similarity estimation and enable more reliable metabolite annotation in complex mixtures.

## Introduction

Mass spectrometry-based metabolomics has emerged as a cornerstone technology in natural products research and drug discovery^1-2^, enabling high-throughput analysis of complex molecular mixtures^3-4^. The field faces significant challenges in compound identification and dereplication, particularly when analyzing structurally diverse natural products^5-10^. Traditional approaches combining nuclear magnetic resonance (NMR) spectroscopy and mass spectrometry (MS) require extensive expertise and are often time-consuming, creating a bottleneck in the discovery pipeline^11-12^. Recent advances in machine learning (ML) have introduced transformative solutions to these challenges, offering unprecedented opportunities for rapid and accurate compound identification^13-17^. The integration of ML with mass spectrometry has catalyzed several innovative approaches. FastEI combines Word2vec spectral embedding with hierarchical navigable small-world graphs (HNSW) to achieve ultra-fast spectrum matching^18-20^. SMART 2.0 employs convolutional neural networks for rapid natural product annotation^21-23^, while MS2Query leverages mass spectral embedding-based chemical similarity predictors for complex mixture analysis^24^. Despite these advances, current methods still face limitations in accurately capturing the complex relationships between spectral similarity and structural similarity, particularly for larger molecules (>800 Da) and structurally complex natural products^25-27^. A critical limitation of existing approaches lies in their reliance on traditional similarity metrics, such as cosine similarity, which often fail to capture the hierarchical nature of mass spectral fragmentation patterns^28-30^. Furthermore, current methods struggle with: limited coverage of spectral libraries, computational inefficiency in large-scale searches, and inability to effectively handle spectral variations across different experimental conditions. These challenges particularly impact natural product research, where structural diversity and complexity demand more sophisticated analytical approaches. Machine learning-based solutions have shown promise in addressing these limitations by learning complex patterns from large datasets^31-33^. However, existing ML approaches often treat mass spectral data as simple numerical vectors, overlooking the inherent sequential and hierarchical nature of fragmentation patterns. Recent developments in natural language processing, particularly transformer-based architectures, offer new possibilities for modeling such complex relationships. These advances suggest that treating mass spectral analysis as a language understanding problem could provide more nuanced and accurate similarity assessments.

Here, we present BertMS, a novel approach that reimagines mass spectral analysis through the lens of natural language processing. By adapting the bidirectional encoder representations from transformers (BERT) architecture to mass spectral data, BertMS learns contextualized representations of fragment ions, enabling more accurate similarity assessments. This approach addresses key limitations of existing methods and is readily scalable to high-throughput metabolomics analysis.

## Results

### Explaining Peak matching with BertMS

The framework of BertMS utilizes a transformer architecture to enhance mass spectral analysis through context-aware fragmentation pattern learning. The input layer processes tandem mass spectra as sequences of peaks, where each peak’s characteristics (m/z values and intensities) are captured through a specialized tokenization scheme. Peak intensities are normalized, and noise is filtered using an adaptive threshold. Each spectrum S is represented as a sequence of peaks P = {p1, p2, …, pn}, where each peak pi contains mass-to-charge ratio (m/z) and intensity information. Unlike traditional approaches that treat peaks as independent entities, BertMS considers the complete fragmentation context through bidirectional attention mechanisms. The core innovation lies in adapting the BERT architecture for mass spectral data interpretation. Input embeddings are constructed as a combination of three components (**Figure 1**): token embeddings representing individual peaks (incorporating both m/z and intensity information), position embeddings encoding the relative location of fragments, and segment embeddings distinguishing between different spectral regions. This embedding strategy allows BertMS to capture both local peak features and global fragmentation patterns. The multi-head attention mechanism in BertMS enables parallel processing of different fragmentation pathways, particularly beneficial for complex natural products where multiple fragmentation routes coexist. During pre-training, BertMS learns from large-scale spectral datasets through masked peak prediction tasks, where 15% of peaks are randomly masked and the model learns to predict their m/z values and intensities based on surrounding spectral context. This self-supervised learning approach allows the model to develop a deep understanding of fragmentation patterns without requiring explicit structural annotations.

**Figure 1.**
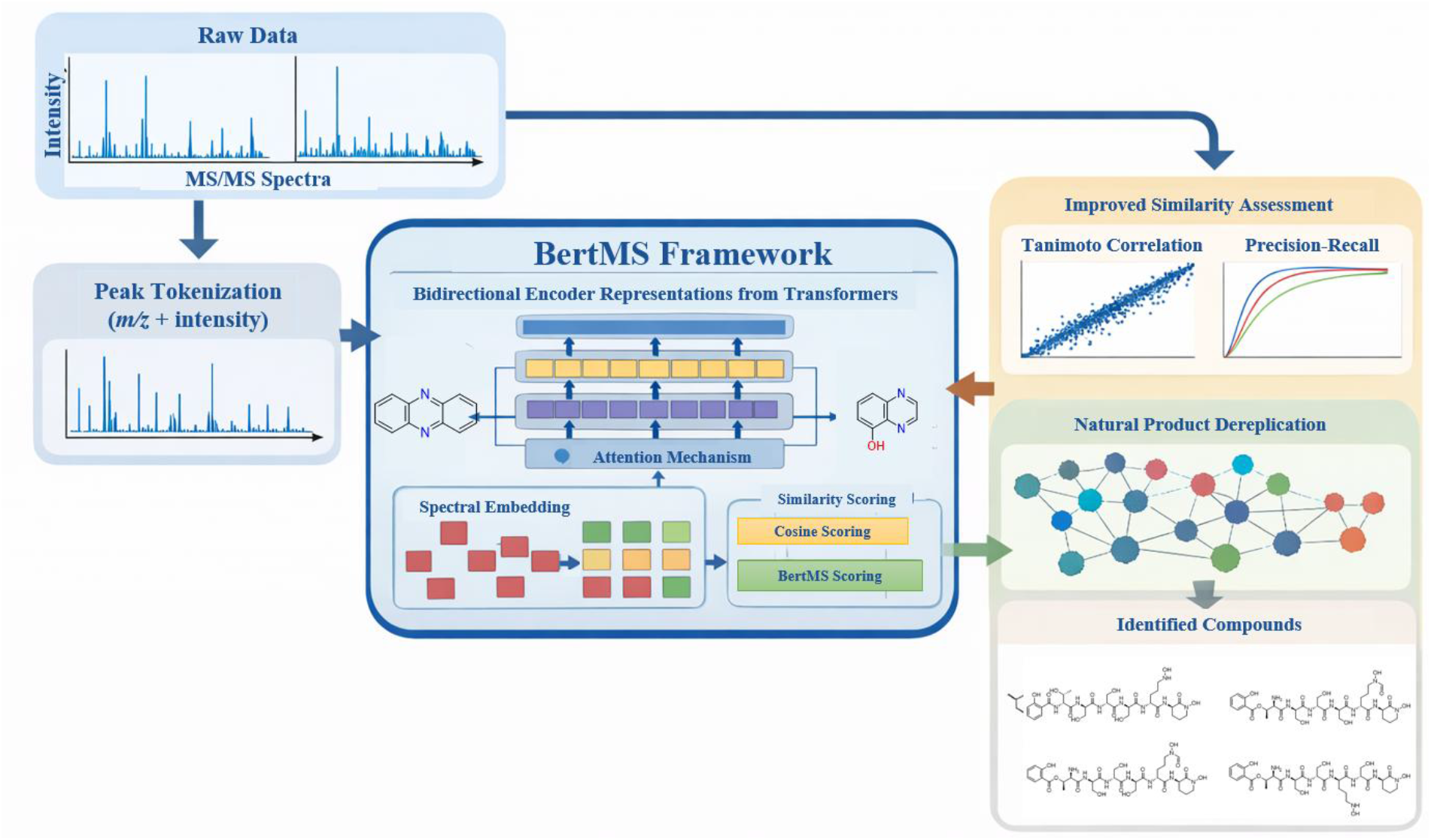
Schematic illustration of the BertMS workflow for spectral similarity analysis. Raw tandem mass spectrometry data are first processed via the MSP platform and transformed into model-compatible inputs. The BertMS core architecture consists of an embedding layer, multi-head self-attention mechanism, feed-forward networks, scaling modules, and SoftMax layers to learn spectral representations. The model produces multiple outputs, including heat-map similarity matrices and node-based relationship networks, facilitating visualization of spectral similarity and compound relationships.

### Spectrum matching performance of BertMS

To evaluate our model’s performance comprehensively, we conducted comparative analyses against established methods including cosine similarity-based approach (GNPS) and word2vec method (Spec2vec). The assessment was performed on a curated dataset comprising 80% of the GNPS library and MoNA tandem mass spectrometry data, with an additional 10% validation set for parameter optimization. The remaining 10% constituted an independent test set for performance evaluation. To ensure unbiased assessment, we carefully eliminated spectral redundancy through InChIKey-based filtering. Through comparative analysis, BertMS shows a relatively stable and approximately linear trend in the trade-off between true and false positives. While Spec2Vec achieves higher initial performance at very low false positive rates (true positive ≈ 0.90 at false positives = 0.005), BertMS exhibits superior scalability and predictability in performance improvement. This linear growth characteristic is particularly evident across the entire false positive range, with BertMS maintaining consistent slope from threshold >0.95 to >0.7. From a quantitative perspective, at a false positive rate of 0.010 per query, BertMS achieves approximately 0.40 true positive rate. Although this is lower than Spec2Vec’s 0.92 and cosine similarity’s rapid initial rise at comparable false positive levels, BertMS’s performance curve shows good linear progression. Notably, as the false positive tolerance increases from 0.018 to 0.038, BertMS’s true positive rate steadily climbs from 0.67 to 0.96, maintaining a nearly constant improvement rate throughout this range. This stable scaling behavior indicates a potential advantage over existing methods. While traditional cosine similarity shows steep but unstable growth concentrated in a narrow false positive range (0.002-0.0035), and Spec2Vec exhibits early saturation, BertMS’s linear behavior suggests better generalization capability. This property becomes particularly advantageous when dealing with complex natural product mixtures or large-scale spectral databases, where controlled false positive rates and predictable performance scaling are critical for practical applications (**Figure 2a**).

**Figure 2.**
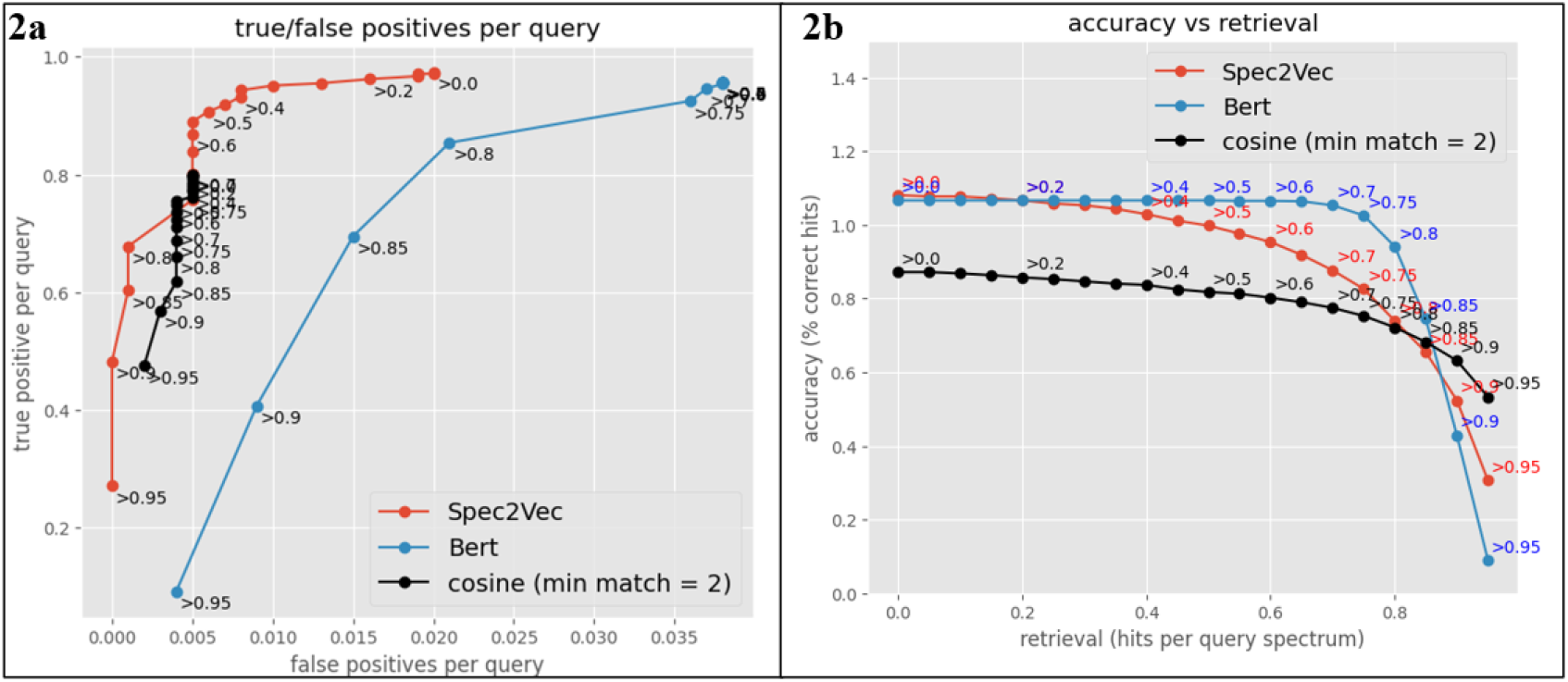
Performance evaluation of spectral matching methods. (2a) Comparison of true vs false positive matches for Spec2Vec, Bert, and cosine similarity approaches across 1000 query spectra. (2b) Trade-off between accuracy and retrieval rate with varying similarity thresholds (>0.0 to >0.95) for each method.

The accuracy evaluation illustrates the performance of three spectral similarity methods across a range of similarity thresholds. Overall, identification precision decreases as the similarity threshold becomes more stringent. BertMS maintains stable and superior performance within the practically relevant similarity range (0.0–0.85). As the threshold increases from 0.0 to approximately 0.75, BertMS consistently preserves high precision. In contrast, Spec2Vec begins to show a noticeable decline at similarity thresholds of approximately 0.4–0.6, whereas conventional cosine similarity exhibits consistently lower precision throughout the evaluated range (typically around 0.85–0.90). Notably, BertMS still achieves approximately 0.80 precision at a similarity threshold of 0.8, demonstrating strong robustness in moderate-to-high similarity regimes. When the similarity threshold exceeds 0.9, the precision of BertMS drops sharply (approximately 0.1–0.2); however, the recall in this region also falls below 10%, making this regime less informative for practical evaluation. Within the operationally meaningful similarity range (threshold ≤0.85), BertMS achieves average precision improvements of approximately 15–20% over Spec2Vec and 20–25% over conventional cosine similarity, highlighting its effectiveness for accurate mass spectral interpretation and compound identification. (**Figure 2b**).

Through quantitative analysis of experimental data, BertMS shows improved performance in molecular similarity assessment tasks. First, from the perspective of performance stability, BertMS maintains a Tanimoto score of approximately 0.42-0.45 over the entire evaluation interval (0-1%). This robust performance is significantly better than other baseline methods. Especially in the key interval of top 0.25%-1%, the fluctuation range of its performance curve is significantly smaller than that of Spec2Vec and cosine similarity methods. Judging from the improvement in algorithm performance, BertMS has achieved about 25% performance improvement compared to the traditional cosine similarity method (about 0.32-0.35). This improvement is consistent across similarity thresholds, indicating robust performance of the model. It is worth noting that even under more stringent filtering conditions (close to the 0.1% area), BertMS still maintains a relatively stable performance advantage. This feature has important practical value in practical applications (**Figure 3**).

**Figure 3.**
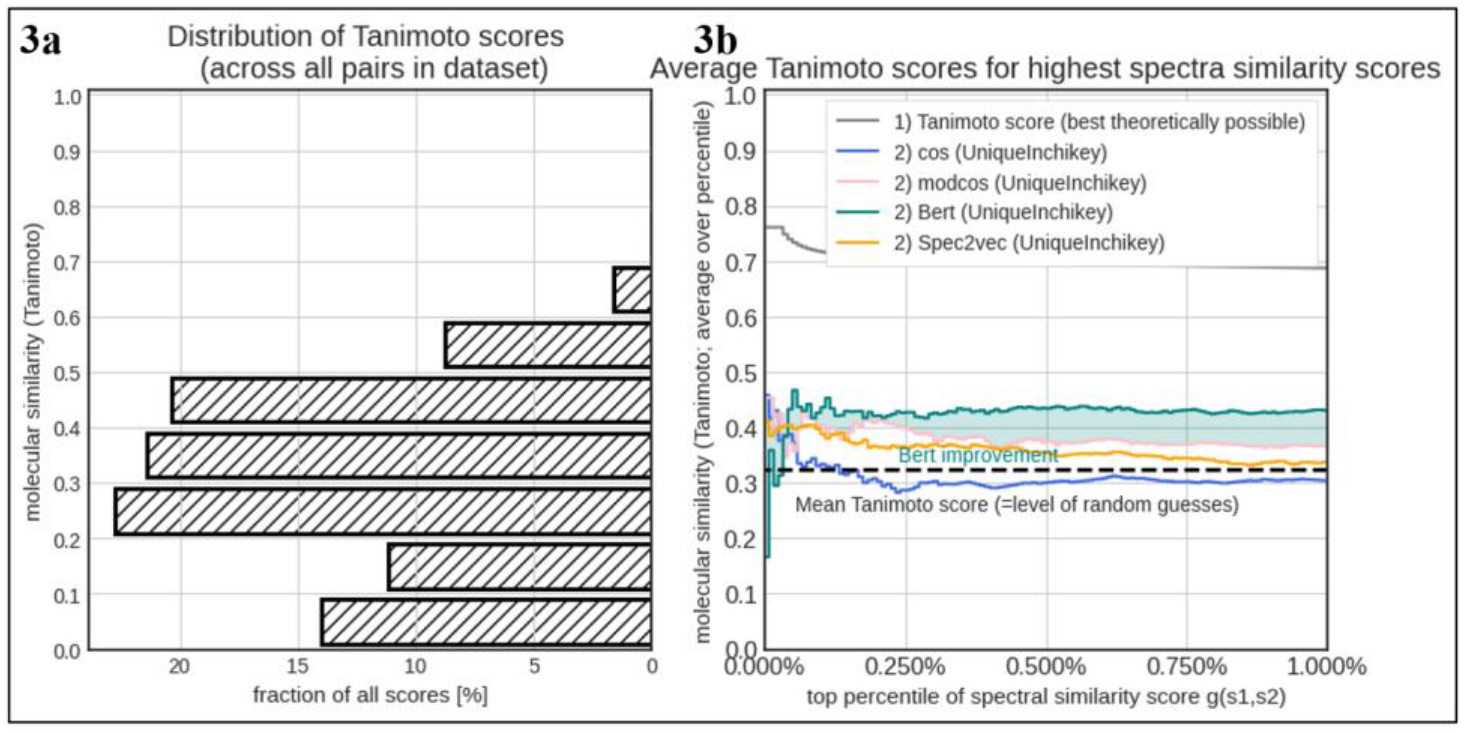
Tanimoto score analysis. (3a) distribution of Tanimoto molecular similarity scores across all spectrum pairs in the dataset. (3b) comparison of average Tanimoto scores for different spectral similarity methods (cosine, modified cosine, BertMS, and Spec2Vec) against the theoretical maximum, showing relative performance across different percentiles of similarity scores.

### Applicability of BertMS to local measured compounds and candidates in library

To validate the model’s capability in learning structural information from spectral fragmentation patterns, we evaluated BertMS performance on experimentally acquired tandem mass spectra from 14 compound pairs (G1-G14) isolated in our laboratory. Figure 4 presents a critical comparative analysis where Tanimoto coefficients, calculated directly from molecular structures, serve as the ground truth labels representing true structural similarity. The spectral similarity scores from BertMS, Spec2Vec, cosine similarity, and modified cosine similarity are evaluated based on their concordance with these structure-based Tanimoto coefficients. Notably, the superior performance of a spectral similarity method is not determined by achieving higher absolute scores, but rather by the degree of agreement with the Tanimoto reference values, indicating successful extraction of structural information from mass spectral data.

**Figure 4.**
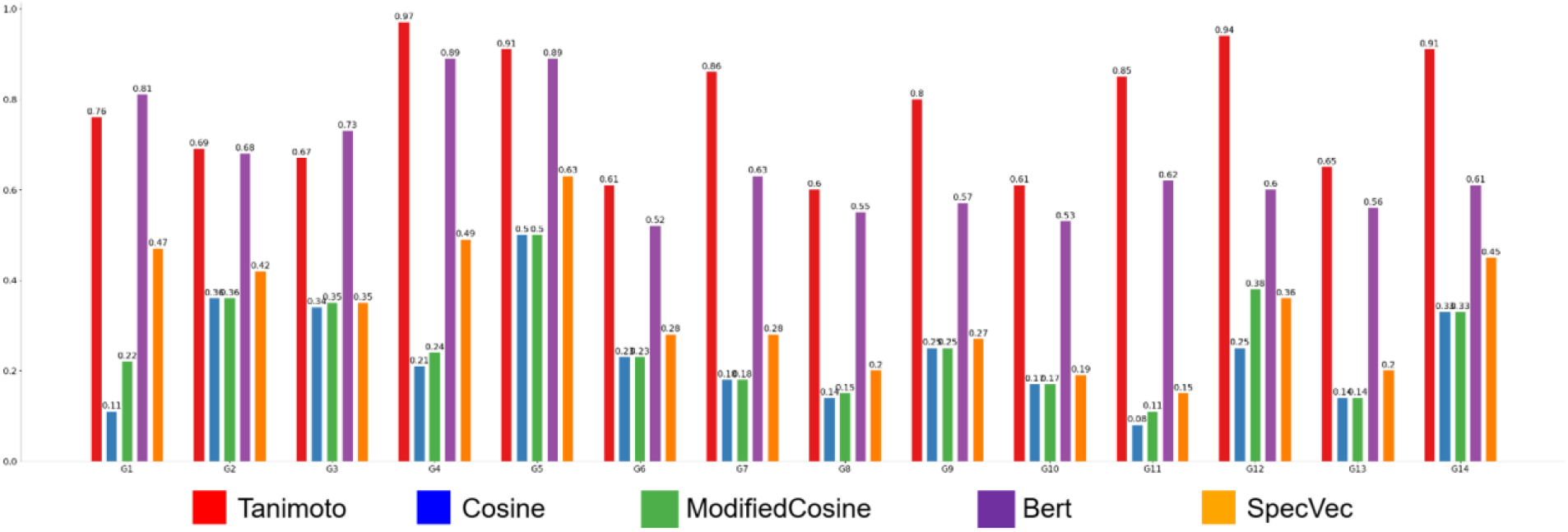
Comparison of different similarity metrics across quarterly periods (G1-G14).

The results demonstrate that BertMS exhibits good agreement with structure-based similarity across most sample groups. For instance, in G1, BertMS achieves 0.81 compared to the Tanimoto reference of 0.76 (deviation: 0.05), while in G4, BertMS scores 0.89 against Tanimoto’s 0.97 (deviation: 0.08). In contrast, conventional cosine-based methods show substantial deviations from the structural ground truth, typically scoring 0.10-0.35 when Tanimoto values range from 0.60-0.97, indicating their failure to capture true structural relationships. Spec2Vec demonstrates intermediate performance with scores of 0.20-0.61, showing improved but still incomplete learning of structural features. This pronounced concordance between BertMS predictions and structure-based Tanimoto coefficients suggests that the transformer-based architecture is able to capture implicit structural representations from fragmentation patterns alone, without direct access to molecular structure information during training. Essentially, BertMS bridges the gap between spectral similarity and structural similarity by learning that similar fragmentation patterns typically arise from structurally related molecules. This capability is particularly valuable in metabolomics workflows where molecular structures are unknown a priori, and spectral data serve as the sole source of structural inference.

A critical technical advantage of BertMS over Spec2Vec lies in its treatment of new spectral peaks. Spec2Vec, based on the Word2Vec architecture, can only generate embeddings for peaks (m/z values) that appeared in its training vocabulary; any unseen peaks encountered in real-world samples are simply ignored during similarity calculation, leading to incomplete spectral representation and potential loss of structurally informative fragments. In contrast, BertMS employs a transformer-based tokenization strategy that generates contextualized embeddings for all spectral peaks, regardless of whether they were observed during training. This design allows BertMS to maintain performance when analyzing compounds or rare fragmentation patterns that were absent from the training data, a scenario frequently encountered in natural product discovery and untargeted metabolomics. The superior concordance between BertMS predictions and structure-based Tanimoto coefficients, particularly for laboratory-isolated compounds that likely contain novel structural features, provides evidence that the model successfully learns generalizable fragmentation-structure relationships rather than merely memorizing training examples. This enhanced generalizability is particularly valuable in metabolomics workflows where the majority of detected features represent unknown or poorly characterized molecules, and the ability to accurately assess structural similarity from spectral data alone—without requiring prior knowledge of specific fragment masses—represents a substantial advancement for structure elucidation and compound dereplication in complex natural product mixtures

We employed the Tanimoto coefficient as the similarity index for compound comparison, selecting a database of compounds for our analysis. Our team conducted empirical validation of the model’s performance using this database. The results demonstrate that in the similarity comparisons of compounds using the Tanimoto index, BertMS consistently yields reasonable similarity scores.

In contrast, BertMS employs a tokenization strategy that allows both previously observed and unseen peaks to be embedded into the model representation space. This design enables BertMS to retain spectral information from new fragment ions rather than discarding them during preprocessing. Consequently, the model exhibits improved robustness and generalization when applied to spectra containing previously unobserved peaks, which are commonly encountered in natural product discovery and untargeted metabolomics studies.

Given these advantages, BertMS is particularly suitable for large-scale spectral analysis scenarios where unknown metabolites are frequently encountered. One important application is molecular networking; an emerging analytical framework widely used for metabolomic data interpretation and natural product discovery. The robust association between BertMS similarity metrics and molecular structural concordance therefore provides a opportunity to explore its application in molecular networking. To evaluate BertMS performance in unknown compound dereplication, we conducted analytical experiments on heterogeneous metabolite datasets derived from microbial sources.

Within the molecular networking framework, individual spectra are represented as nodes, and edges are established when similarity scores exceed predefined thresholds. Most existing approaches rely on modified cosine similarity to determine these connections. The resulting network structures enable the identification of molecular families through cluster analysis and facilitate the dereplication of structurally related metabolites.

### Unknown compounds dereplication application

The robust association between BertMS similarity metrics and molecular structural concordance presented a opportunity to investigate its implementation in molecular networking - an innovative analytical framework for metabolomic data interpretation. To assess BertMS’s efficacy in novel compound dereplication, we executed extensive analytical protocols on heterogeneous metabolite assemblages derived from microbial origins. In molecular networking, each spectrum is treated as a node, and edges are defined when similarity scores exceed a given threshold. Contemporary methodologies predominantly employ modified cosine calculations for connection determination.

These networked architectures facilitate the elucidation of molecular families through systematic cluster analysis.

In this investigation, we demonstrate an application of this cheminformatic approach for the automated characterization of complex natural products from cyanobacterial extract matrices. The microbial strain HDN19-252, taxonomically classified as *Nocardiopsis aegyptia*, was obtained from Antarctic samples. Following rice-based fermentation, metabolite extraction was performed using methanol, and the resultant extract underwent LC-MS/MS analysis utilizing UPLC instrumentation. The acquired data underwent preprocessing and subsequent analysis via the BertMS computational platform, with network visualization accomplished through Cytoscape software. Molecular network analysis revealed three distinctive clusters through systematic evaluation. Subsequent isolation and structural characterization of the compounds yielded a new class of polypeptides (**Figure 6**), designated as nocaslide A-F (**1-6**), representing a previously undescribed class of siderophores, alongside a new neurotensin antagonist peptide analog, neuroslide A (**7**). This report introduces an extensively enhanced, user-friendly iteration of MS/MS analysis platform, demonstrating its effectiveness in rapid natural product structure classification.

**Figure 6.**
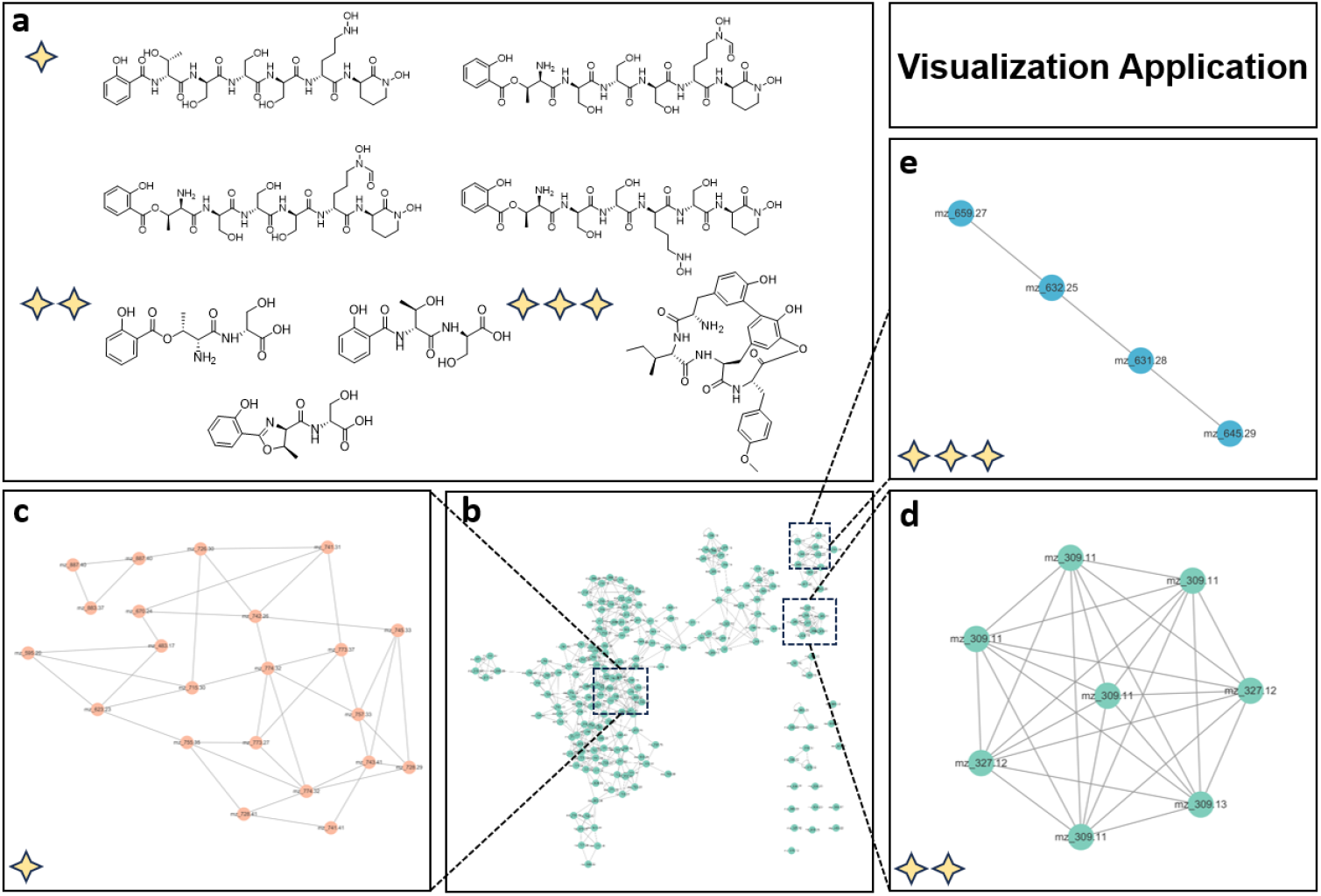
The application of BertMS on microbial metabolites.

**Figure 7:**
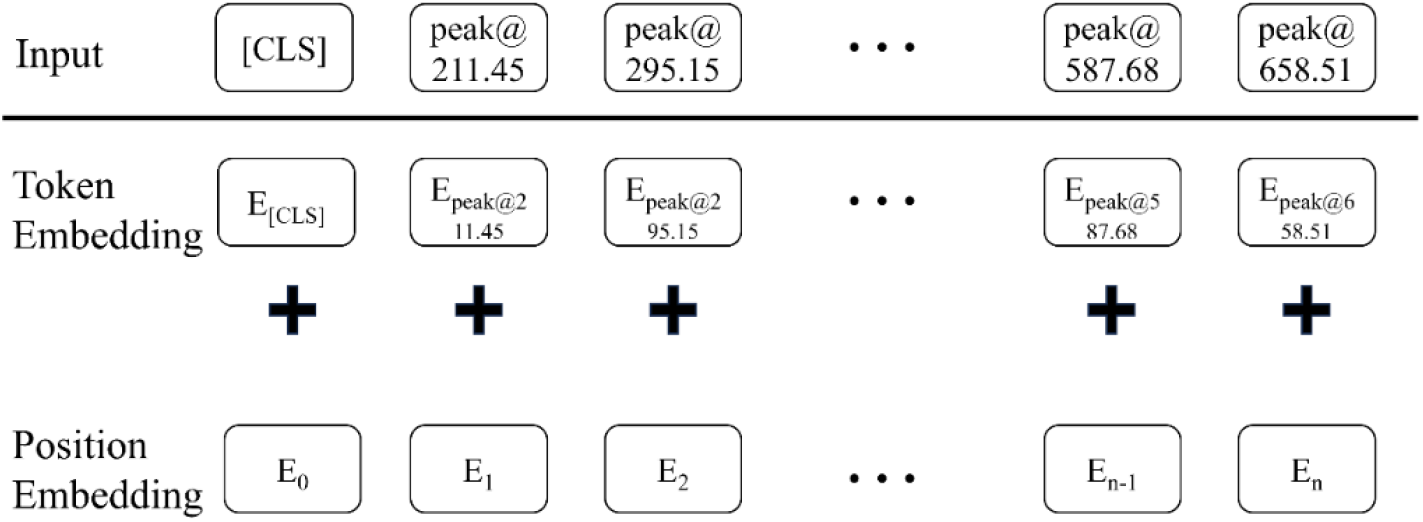
BertMS Input Representation: The input embeddings for BertMS are the sum of the token embeddings and position embeddings.

## Discussion

In this study, we introduce BertMS, a transformer-based framework for spectral similarity estimation in tandem mass spectrometry. By modeling spectra as sequences of fragment peaks and leveraging bidirectional attention mechanisms, the method provides a contextual representation of spectral features. Our results indicate that BertMS improves the consistency between spectral similarity and structural similarity compared with commonly used approaches such as cosine similarity and Spec2Vec. This improvement is particularly evident in molecular similarity assessment and in applications involving complex mixtures. An important advantage of BertMS is its ability to represent previously unseen peaks, which may contribute to improved generalization when analyzing novel compounds. This property is relevant for metabolomics applications, where unknown or poorly characterized metabolites are frequently encountered. Despite these advantages, several limitations should be considered. As a data-driven approach, the performance of BertMS depends on the diversity and coverage of the training dataset. In addition, while the model captures relationships between spectral features, further work is needed to improve interpretability and to better understand how these representations relate to chemical structure. BertMS complements existing spectral similarity methods by improving consistency between spectral and structural similarity. Its computational efficiency and scalability make it suitable for large-scale applications, including library searching and molecular networking. Future work may explore integration with other computational tools and extension to additional types of mass spectrometry data.

## Methods

### Problem description

#### Algorithm 1

MS-BERT Prc-Lraining Framework

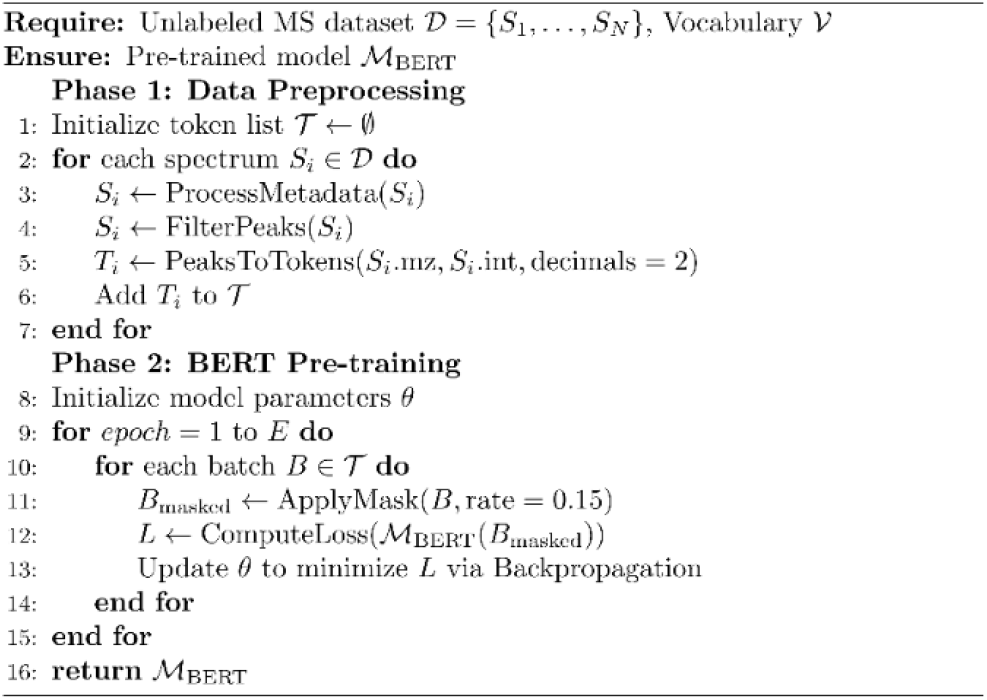

#### Algorithm 2

BERT-MN: Network Construction and Clustering

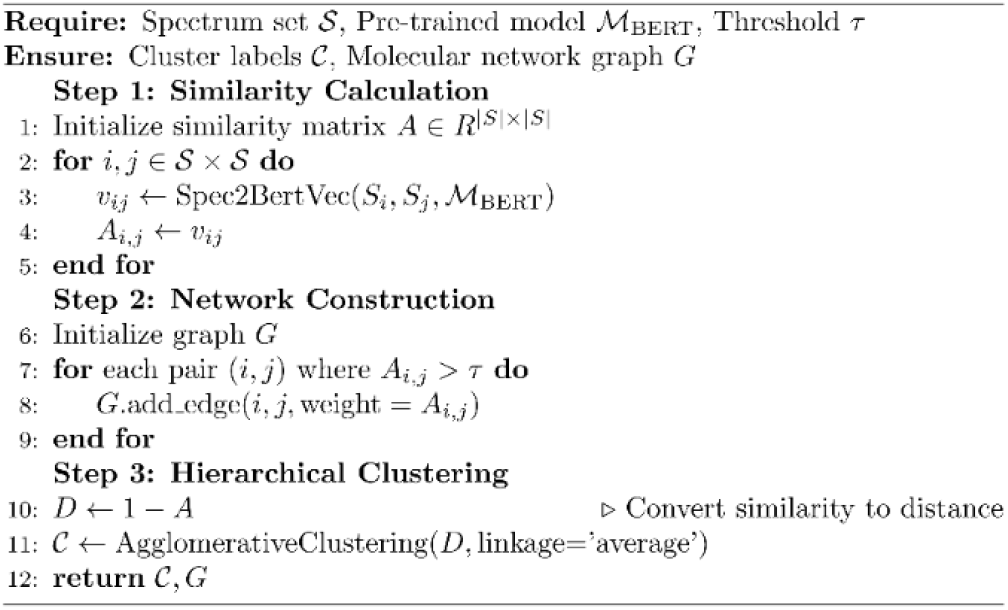

The interpretation of mass spectral data for compound identification presents unique challenges analogous to natural language understanding (NLU) tasks. In this work, we formulate mass spectral analysis as a representation learning problem, where each spectrum is mapped into a high-dimensional embedding space that preserves both local fragment relationships and global structural patterns. To overcome the limitations of static embedding models like Word2Vec in handling unseen tokens (or rare fragments), we employ BERT as the spectral encoding backbone. While the standard BERT framework comprises both pre-training and fine-tuning phases, this study specifically leverages the pre-training stage to generate unsupervised $d$-dimensional representations of mass spectra, serving as a robust feature extractor for downstream tasks.

### Network Architecture and Problem Formulation

The BertMS model employs a multi-layer bidirectional Transformer encoder architecture, leveraging bidirectional self-attention to capture contextual dependencies across the entire input sequence. Unlike Natural Language Processing (NLP) tasks where sentence pairs are often used, we treat each mass spectrum as a single coherent sequence given the specific characteristics of mass spectrometry (MS) data. Formally, let a mass spectrum *S* be represented as a set of *n* peaks *S* = (*m*/*z*_1_, *I*_1_), (*m*/*z*_2_, *I*_2_), …, (*m*/*z*_*n*_, *I*_*n*_) where *m*/*z* denotes the mass-to-charge ratio and *I* denotes intensity. Our objective is to learn a mapping function *f*: *S* → *R*^*d*^ that transforms the discrete spectral data into a continuous *d*-dimensional embedding space. Within this space, the structural similarity between two spectra can be quantified by the euclidean or cosine distance between their corresponding embeddings. To effectively process MS data, we design a hierarchical feature representation: **Peak-level encoding:** Each peak (*m*/*z, I*) is discretized and projected into a token embedding space. **Position-aware attention**: Relative positions of spectral fragments are preserved via learnable positional embeddings, allowing the model to distinguish between low-mass and high-mass regions. **Global context modeling**: A special [CLS] token is prepended to the sequence to aggregate whole-spectrum information for downstream classification or retrieval tasks.\end{itemize}. The core of the encoder is the Multi-head Self-attention mechanism. For a given input, the attention head is computed as: “*Head*”_*i*_ = “*Attention*”(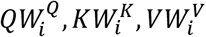) where Q, K, V are linear transformations of the input embeddings. The final representation *H* is obtained by concatenating all heads: *H* = \*text*{*Concat*}(\ *text*{*head*}_1_,\*dots*,\*text*{*head*}_*h*_)*W*^*0*^.

### Pre-training Strategy and Objectives

Given the unlabeled nature of large-scale MS databases, we adopt a self-supervised learning approach. Specifically, we utilize the Masked Language Model (MLM) task to enable the model to learn complex fragmentation patterns and peak correlations. The pre-training objective consists of three key aspects: (1) Masked Peak Prediction, where the model reconstructs missing peaks based on context; (2) Intensity Prediction, enabling the model to learn abundance relationships; and (3) Fragment Correlation Learning, which captures implicit chemical fragmentation rules. Following the standard BERT protocol, we randomly mask 15% of the tokens in each spectrum. The masking strategy is implemented as follows: 80% of the selected tokens are replaced with a special [MASK] token. 10% are replaced with a random token from the vocabulary. 10% remain unchanged to preserve real data distribution. The model predicts the original tokens from the masked input using a SoftMax layer over the vocabulary, optimized via cross-entropy loss. This forces MS-BERT to capture both local peak interactions and global spectral features, generating chemically meaningful embeddings.

#### Data Preparation

Firstly, this study utilizes a mass spectrometry dataset (LS-MS) comprising 154,820 spectra, which is provided by GNPS (https://gnps-external.ucsd.edu/gnpslibrary/ALL_GNPS.json). The raw data is now accessible at https://doi.org/10.5281/zenodo.3979010. Metadata was cleaned and corrected using matchms^34^, resulting in 94,121 spectra annotated with InChIKey. Similar to Spec2vec^35^, we employed key metadata information (including compound names, chemical formulas, and estimated parent masses) for extensive automated lookup searches on PubChem^36^ via pubchempy^37^, in order to obtain the InChIKey for unlabeled spectra. Additionally, we incorporated mass spectrometry data from our laboratory experiments (details to be supplemented) to test the generalizability of our model.

#### Conversion of Mass Spectra to Documents

To ensure that the underlying Bert^38^ model is trained using documents of uniform size, this is achieved by eliminating lower-intensity, non-informative fragment peaks. Specifically, the maximum number of peaks retained for each spectrum is directly proportional to the estimated parent mass:

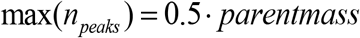

Thus, each mass spectrum is converted into a document. Each peak in the spectrum is represented as a word in the format (rounded to two decimal places). For example, a peak at 211.452 would be converted to “peak@211.45“. The list of words representing all the peaks in a spectrum constitutes the document for that spectrum. Furthermore, the entire vocabulary of words from all spectrum documents is used as the word list during model training. This ensures that words are randomly masked and replaced from this vocabulary during subsequent training of the model.

#### Similarity Score Calculation

The similarity score is derived from the previously trained BertMS model. The BertMS model learns the relationships between words through the Masked Language Model (MLM) task. The hidden word vectors (the mapping of input tokens in the d-dimensional space) are used as word embeddings.

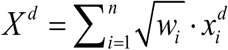

where *w*_*i*_ represents the intensity of the i peak (with all peak intensities normalized to a maximum intensity of 1), and *w*_*i*_ is the hidden vector of the i peak in the d-dimensional space.

The similarity between two mass spectra is calculated using the cosine similarity between their embeddings:

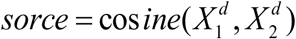

Similarly, to better compare the calculation of BertMS similarity scores, we also computed the Spec2vec, cosine scores, and cosine-corrected scores (implemented using the Python package matchms). For both cosine scores and corrected cosine scores, each peak can only be matched once.

#### Structural Similarity

Currently, most studies consider the Tanimoto similarity score to be one of the most effective methods for assessing the structural similarity between two compounds^39^. Therefore, in this study, the structural similarity corresponding to the two mass spectra data is measured by calculating the Tanimoto similarity (Jaccard index). Specifically, we first compute molecular fingerprints using rdkit^40^ for the molecular structures, and then calculate the Tanimoto similarity between the molecular fingerprints using methods from matchms.

#### Implementation Details

The BertMS model is implemented in PyTorch^41^. The dimension of each word embedding is set to 768. We also adapted the code from Spec2vec and matchms to fit the similarity calculations in this study. For specific hyperparameter settings such as learning rate, weight decay, batch size, and network architecture parameters (number of hidden layers and dimensions), please refer to the code repository.

## Data Availability Code Availability

Data and code will be made publicly available upon reasonable request and will be released through a public repository after peer-reviewed publication.

## Contributions

S.W. designed BertMS algorithm and L.Z. optimized the algorithm. J.P., W.W and X.H. performed the analysis and isolation. L.Z. and S.W. were responsible for manuscript writing. D.L. designed and directed the work. ‡ L.Z. and S.W. contributed equally.

## Acknowledgement

This work is supported by the Fundamental Research Funds for the Central Universities (202372009, and 202572006), the National Key R&D Program of China (2024YFC2816004), the Qingdao Marine Science and Technology Center (2022QNLM030003-1), the Key R&D Program of Hainan Province (ZDYF2023SHFZ144), and the Taishan Scholar Distinguished Expert Program in Shandong Province (tstp20240504).

