## Supplementary Information for "BertMS-enabled molecular networking for unknown compounds dereplication"

#### Author Information

Luning Zhou‡, Shuang Wu‡, Jixing Peng, Xiaofei Huang, Wenxue Wang and Dehai Li*

#### Corresponding author

#### Experimental section

##### General experimental procedures

UV spectra were recorded on HITACHI 5430 spectrophotometer (Hitachi Ltd., Tokyo, Japan). LC-MS/MS data were obtained on AB SCIEX ZENO Q-TOF 7600NMR. spectra were recorded on a Bruker AVANCE NEO 400 MHz spectrometer, an Agilent DD2-500 spectrometer (Agilent Technologies, Palo Alto CA., America) and a JEOL JNM-ECP600 spectrometer (JEOL (BEIJING) Co., Ltd. Shanghai Branch). HRESIMS data were obtained using a Thermo Scientific LTQ Orbitrap XL mass spectrometer (Thermo Fisher Scientific, Waltham, MA, America). Column chromatography (CC) was performed with silica gel (Qingdao Marine Chemical Factory, Qingdao, China). The compounds were purified by a HITACHI 1110 system (Hitachi Ltd., Tokyo, Japan) equipped with a 1430 PDA detector and a C18 column (YMC-Pack ODS-A, 10 × 250 mm, 5 $\mu$m, 3 mL/min).

**Table 1**. LC-MS/MS analysis method

| Time | A (water/formic acid) % | B (MeCN) % | Flow |
|---|---|---|---|
| 0 | 90 | 10 | 0.3 |
| 1 | 90 | 10 | 0.3 |
| 7 | 0 | 100 | 0.3 |
| 10 | 0 | 100 | 0.3 |
| 10.1 | 90 | 10 | 0.3 |

##### The extraction and isolation of *Nocardiopsis aegyptia* HDN19-252

The strain *Nocardiopsis aegyptia* HDN19-252 (GenBank accession number MN822699) was isolated from zoological specimens obtained from Antarctic coordinates (61°42′28″ S, 57°38′22″ W). The strain has been deposited and maintained in the repository of the Key Laboratory of Marine Drugs, housed within the School of Medicine and Pharmacy at Ocean University of China, situated in Qingdao, People's Republic of China. *Nocardiopsis aegyptia* HDN19-252 was cultured in 1 L Erlenmeyer flasks containing 200 g of culture medium composed of 80 g of rice and 120 g of seawater, pH = 7.0 (in seawater collected from Huiquan Bay, Yellow Sea) at 28 °C for

30 days on stable fer-mentation. A total of 70 bottles of the culture medium were extracted with MeOH (3× 20 L) to generate a crude extract (50.5 g).

LC-MS/MS analysis was performed using a UHPLC system combined with a hybrid Quadrupole-Orbitrap mass spectrometer (AB SCIEX ZENO Q-TOF 7600). As a mobile phase, 0.1% formic acid in $H_2O$ (A) and HPLC-grade MeCN (B) were used in negative-ionization conditions. The elution gradient conditions of LC-MS/MS were as follows, based on times (t): t = 0−1 min, hold at 10% B; t = 1−7 min, increased to 100% B linearly; t = 7−10 min, thold at 100% B; t =10−10.1 min, returned to initial conditions and hold at 10% B to re-equilibrate the column. The elution velocity and injection volume were 0.3 mL/min and 1 μL, respectively. All MS/MS data were converted to msp format files by MSConvert software (Ver. 3.0.20169, MsConvert, ProteoWizard) and MS-finder (Ver. 3.6.1). The .xgmml spectral network files were visualized through Cytoscape (Ver. 3.8.0, Cytoscape, NRNB.)

The crude extract was separated by ODS column and eluted with mixtures of MeOH/$H_2O$ (20%, 30%, 40%, 50%, 100%) to give five fractions. Each fraction was divided to four subfractions. LCMS were used to detected the target compound peak by molecular weight. Fr.2.1 was purified by semi-preparative HPLC to obtain **1** (10.1 mg, $t_R$ = 15.1 min) and **2** (7.5 mg, $t_R$ = 15.3 min). Fr.2.3 was purified by semi-preparative HPLC to afford **3** (6.5 mg, $t_R$ = 14.2 min) and **4** (5.3 mg, $t_R$ = 15.0 min). Fr.3.1 was purified by semi-preparative HPLC to afford **5** (4.2 mg, $t_R$ = 13.0 min), **6** (4.5 mg, $t_R$ = 13.2 min) and **7** (3.5 mg, $t_R$ = 21.0 min). Fr.3.2 was purified by semi-preparative HPLC using a stepped gradient elution to obtain **8** (6.5 mg, $t_R$ = 19 min).

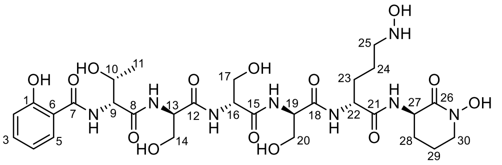

1

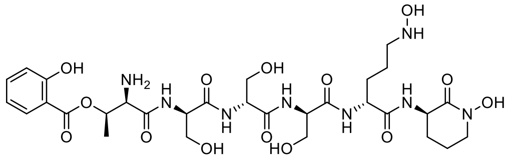

2

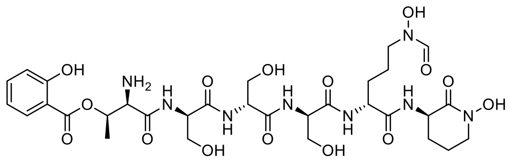

3

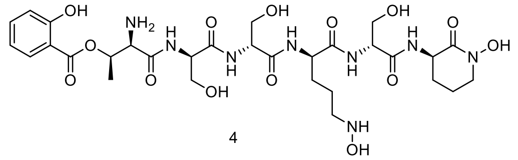

4

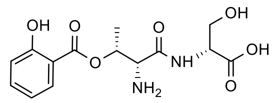

5

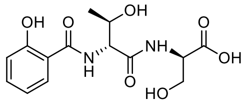

6

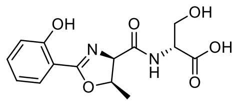

7

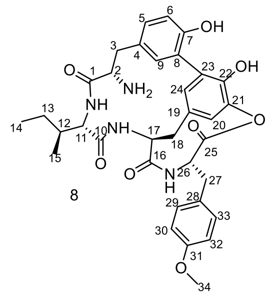

8

The molecular formula of **1** was established as $C_{30}H_{46}N_8O_{14}$ on the basis of HRESIMS data. Analysis of $^1$H, $^{13}$C, and HSQC NMR data indicated the presence of thirty carbons, including seven carbonyls, six aromatic carbons with for being protonated, eleven methines, nine methylenes, one methyl groups, and nine exchangeable protons. Analyses of COSY and TOCSY data reveled six amino acid spin systems consistent with a threonine derived moiety, three serine derived moiety, and two modified ornithine moieties. The coupling constants of H-2 (d, $J$ = 8.1 Hz), H-3 (t, $J$ = 7.6 Hz), H-4 (t, $J$ = 7.6 Hz) and H-5 (d, $J$ = 8.1 Hz) and COSY correlation of H-2/H-3/H-4/H-5, together with the HMBC correlations from H-5 to C-1, C-6 and C-7 indicated the presence of salicylic acid substructure. The amino acid sequence was assigned using a combination of HMBC, NOESY and TOCSY correlations, and MS/MS experiments. The first conjunction between salicylic acid substructure and threonine were supported by HMBC correlations from NH-9 to C-7 and C-9, as well as TOCSY correlation of NH-9/H-9/H-10/H-11. Furthermore, HMBC correlation from NH-13 to C-8 and C-13 and the TOCSY correlation of NH-13/H-13/H-14 indicated that serine-1 was attached to C-8 through N-13. HMBC correlation from NH-16 to C-12 and C-16 and NH-19 to C-15 and C-19, as well the TOCSY correlation of NH-16/H-16/H-17 and NH-19/H-

19/H-20 unveil that the connectivity between serine-1-serine-2-serine-3. The linkage between serine-3 and modified ornithine-1 moieties were determined by the HMBC correlation from NH-22 to C-18 and C-22, together with the TOCSY of NH-22/H-22/H-23/H-24/H-25. The last ring system of modified ornithine-2 and the conjunction between modified ornithine-1 and ornithine-2 was established by the HMBC correlation from NH-27 to C-21 and C-27, as well as the TOCSY of NH-27/H-27/H-28/H-29/H-30. Accordingly, the planar structure of 1 was established. To determine the absolute configuration of **1**, Marfey's methods were used and the final absolute configuration of **1** was deduced to be 9*R*, 10*R*, 13*R*, 16*R*, 19*R*, 22*R*, 27*R*.

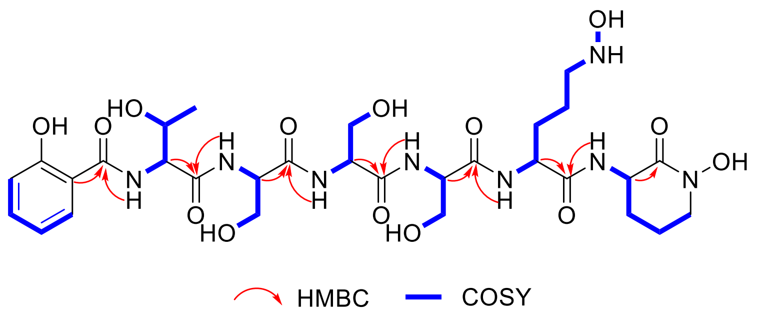


**Table 1**. $^{1}$H NMR (400 MHz) and $^{13}$C NMR (150 MHz) of **1** in DMSO-$d_6$ ($\delta$ in ppm, $J$ in Hz)

| **1**Unit | Position | $\delta_C$ type | $\delta_H$ ($J$ in Hz) |
|---|---|---|---|
| SA | 1 | 158.2, C | |
| | 2 | 130.4, CH | 7.91, d (8.3) |
| | 3 | 133.9, CH | 7.39, t (7.6) |
| | 4 | 119.7, CH | 6.91, t (7.8) |
| | 5 | 117.5, CH | 6.94, d (7.9) |
| | 6 | 117.9, C | |
| | 7 | 167.1, C | |
| Thr-1 | 8 | 170.9, C | |
| | 9 | 59.8, CH | 4.42, m |
| | 10 | 67.1, CH | 4.07, m |
| | 11 | 20.4, $CH_3$ | 1.11, d (6.4) |
| | 9-NH | | 8.87, d (6.9) |
| Ser-1 | 12 | 170.7, C | |
| | 13 | 55.8, CH | 4.32, m |
| | 14 | 62.1, $CH_2$ | 3.62, m |
| | 13-NH | | 8.24, d (7.4) |
| Ser-2 | 15 | 170.6, C | |
| | 16 | 55.7, CH | 4.32, m |

| | | | |
|---|---|---|---|
| | 17 | 62.1, $CH_2$ | 3.62, m |
| | 16-NH | | 8.04, d (7.4) |
| Ser-3 | 18 | 171.2, C | |
| | 19 | 55.7, CH | 4.32, m |
| | 20 | 62.1, $CH_2$ | 3.62, m |
| | 19-NH | | 8.00, d (7.3) |
| Haorn-1 | 21 | 170.3, C | |
| | 22 | 52.2, CH | 4.30, m |
| | 23 | 28.0, $CH_2$ | 1.88, 1.65, m |
| | 24 | 20.8, $CH_2$ | 1.88, m |
| | 25 | 51.7, $CH_2$ | 3.47, m |
| | 22-NH | | 8.17, d (8.4) |
| Haorn-2 | 26 | 165.2, C | |
| | 27 | 50.1, CH | 4.29, m |
| | 28 | 29.7, $CH_2$ | 1.79, 1.58, m |
| | 29 | 20.0, $CH_2$ | 1.63, m |
| | 30 | 50.5, $CH_2$ | 3.08, m |
| | 27-NH | | 7.96, d (8.4) |

Compound **2**, obtained as amorphous oil, has a molecular formula of $C_{30}H_{46}N_8O_{14}$ deduced by the HRESIMS. Analysis of the 1D and TOCSY NMR data deduced that it was same as **1**. The only difference was the conjunction between salicylic acid substructure and threonine moiety, which was that O-10 was attached to C-7 through an ester bond, supported by the key HMBC correlation from H-10 to C-7 and the hydrogen integral of $NH_2$-9 was 2. Hence, the planar structure of **2** was determined and the absolute configuration were established as 9*R*, 10*R*, 13*R*, 16*R*, 19*R*, 22*R*, 27*R*.

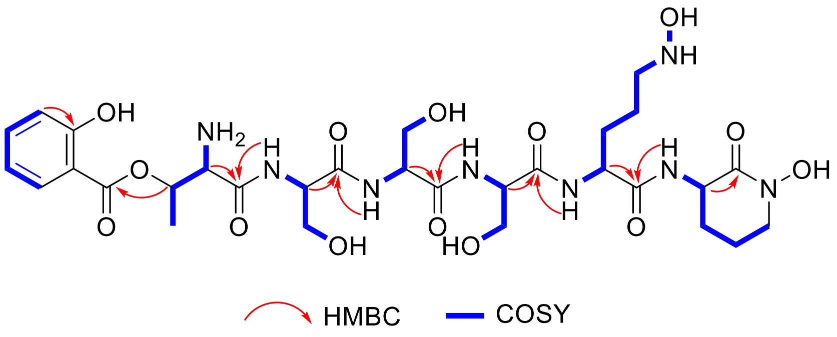


**Table 2.** $^1$H NMR (400 MHz) and $^{13}$C NMR (125 MHz) of **2** in DMSO-$d_6$ ($\delta$ in ppm, $J$ in Hz)

| **2** Unit | Position | $\delta_C$ type | $\delta_H$ ($J$ in Hz) |
|---|---|---|---|

| | | | |
|---|---|---|---|
| SA | 1 | 160.7, C | |
| | 2 | 131.7, CH | 7.97, d (8.3) |
| | 3 | 136.4, CH | 7.52, t (7.6) |
| | 4 | 119.7, CH | 6.93, t (7.8) |
| | 5 | 117.7, CH | 6.97, d (7.9) |
| | 6 | 113.4, C | |
| | 7 | 167.9, C | |
| Thr-1 | 8 | 166.2, C | |
| | 9 | 55.3, CH | 4.25, m |
| | 10 | 70.4, CH | 5.43, m |
| | 11 | 16.8, $CH_3$ | 1.41, d (6.4) |
| | 9-NH | | 8.46, m |
| Ser-1 | 12 | 169.7, C | |
| | 13 | 55.4, CH | 4.54, dd (7.7) |
| | 14 | 62.3, $CH_2$ | 3.53, m |
| | 13-NH | | 8.89, d (8.1) |
| Ser-2 | 15 | 170.4, C | |
| | 16 | 55.5, CH | 4.37, m |
| | 17 | 62.3, $CH_2$ | 3.57, m |
| | 16-NH | | 8.22, m |
| Ser-3 | 18 | 171.2, C | |
| | 19 | 55.5, CH | 4.33, m |
| | 20 | 62.4, $CH_2$ | 3.56, m |
| | 19-NH | | 8.01, m |
| Haorn-1 | 21 | 170.2, C | |
| | 22 | 50.0, CH | 4.30, m |
| | 23 | 28.0, $CH_2$ | 1.88, 1.65, m |
| | 24 | 20.8, $CH_2$ | 1.88, m |
| | 25 | 51.7, $CH_2$ | 3.47, m |
| | 22-NH | | 8.23, m |
| Haorn-2 | 26 | 165.2, C | |
| | 27 | 55.7, CH | 4.28, m |
| | 28 | 29.8, $CH_2$ | 1.78, 1.58, m |
| | 29 | 20.1, $CH_2$ | 1.64, m |
| | 30 | 50.4, $CH_2$ | 3.08, m |
| | 27-NH | | 8.09, d (8.0) |

Compound **3,** obtained as amorphous oil, has a molecular formula of $C_{31}H_{46}N_8O_{15}$ deduced by the HRESIMS. Analysis of the 1D and TOCSY NMR data. It was same as **2**. The only difference was an extra N-formyl group in **3**, which supported by the key

HMBC correlation from H-25 to C-26. Similarly, the absolute configuration of compound **3** was finally determined to be 9*R*, 10*R*, 13*R*, 16*R*, 19*R*, 22*R*, 28*R*.

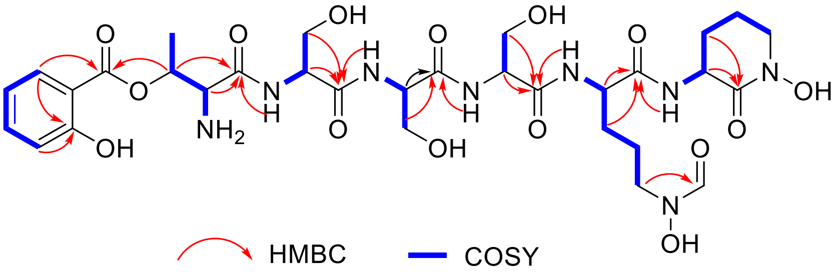


**Table 3**. $^{1}$H NMR (400 MHz) and $^{13}$C NMR (125 MHz) of **3** in DMSO-$d_6$ ($\delta$ in ppm, $J$ in Hz)

| **3** Unit | Position | $\delta_C$ type | $\delta_H$ ($J$ in Hz) |
|---|---|---|---|
| SA | 1 | 160.7, C | |
| | 2 | 131.6, CH | 8.00, d (8.3) |
| | 3 | 136.4, CH | 7.55, t (7.6) |
| | 4 | 119.7, CH | 6.96, t (7.8) |
| | 5 | 117.7, CH | 6.99, d (7.9) |
| | 6 | 113.3, C | |
| | 7 | 168.0, C | |
| Thr-1 | 8 | 166.2, C | |
| | 9 | 55.5, CH | 4.26, m |
| | 10 | 70.4, CH | 5.44, m |
| | 11 | 16.8, $CH_3$ | 1.38, d (6.4) |
| | 9-NH | | 8.48, d (8.2) |
| Ser-1 | 12 | 169.7, C | |
| | 13 | 55.4, CH | 4.51, m |
| | 14 | 62.3, $CH_2$ | 3.55, m |
| | 13-NH | | 8.92, d (8.2) |
| Ser-2 | 15 | 170.4, C | |
| | 16 | 55.5, CH | 4.38, m |
| | 17 | 62.3, $CH_2$ | 3.55, m |
| | 16-NH | | 8.25, br |
| Ser-3 | 18 | 171.5, C | |
| | 19 | 55.5, CH | 4.31, m |
| | 20 | 62.4, $CH_2$ | 3.55, m |
| | 19-NH | | 8.02, m |
| Haorn-1 | 21 | 170.1, C | |
| | 22 | 55.4, CH | 4.25, m |
| | 23 | 29.3, $CH_2$ | 1.68 1.49, m |
| | 24 | 23.4, $CH_2$ | 1.63, m |
| | 25 | 51.6, $CH_2$ | 3.48, m |

| | | | |
|---|---|---|---|
| | 26 | 157.6, CH | 7.9, s |
| | 22-NH | | 8.04, br |
| Haorn-2 | 27 | 165.1, C | |
| | 28 | 49.9, CH | 4.30, m |
| | 29 | 28.0, $CH_2$ | 1.86, 1.67, m |
| | 30 | 20.8, $CH_2$ | 1.90, m |
| | 31 | 51.7, $CH_2$ | 3.46, br |
| | 28-NH | | 8.23, m |

Compound **4,** obtained as amorphous oil, has a molecular formula of $C_{31}H_{46}N_8O_{15}$ deduced by the HRESIMS. Comparison of NMR data revealed that compounds **2** and **4** have certain structural similarities. There were differences in the amino acid positions in the structure and it was determined by HMBC and TOCSY correlations. Finally, the planar structure of **4** was determined and the absolute configuration of 4 was deduced as 9*R*, 10*R*, 13*R*, 16*R*, 19*R*, 24*R*, 27*R*.

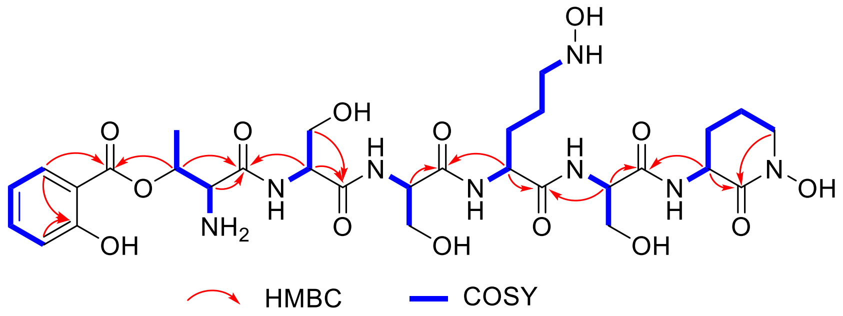


**Table 4**. [1]H NMR (500 MHz) and [13]C NMR (150 MHz) of **4** in DMSO-$d_6$ ($\delta$ in ppm, $J$ in Hz)

| **4** Unit | Position | $\delta_C$ type | $\delta_H$ ($J$ in Hz) |
|---|---|---|---|
| SA | 1 | 160.7, C | |
| | 2 | 131.7, CH | 7.97, d (8.3) |
| | 3 | 136.4, CH | 7.52, t (7.6) |
| | 4 | 119.7, CH | 6.93, t (7.8) |
| | 5 | 117.7, CH | 6.97, d (7.9) |
| | 6 | 113.4, C | |
| | 7 | 168.0, C | |
| Thr-1 | 8 | 166.3, C | |
| | 9 | 55.6, CH | 4.25, m |
| | 10 | 70.5, CH | 5.42, m |
| | 11 | 16.8, $CH_3$ | 1.38, d (6.4) |
| | 9-NH | | 8.48, d (8.2) |
| Ser-1 | 12 | 169.8, C | |
| | 13 | 55.4, CH | 4.51, dd (7.7) |

| | | | |
|---|---|---|---|
| | 14 | 62.3, $CH_2$ | 3.52, m |
| | 13-NH | | 8.90, d (8.2) |
| Ser-2 | 15 | 170.2, C | |
| | 16 | 55.7, CH | 4.38, m |
| | 17 | 62.4, $CH_2$ | 3.55, m |
| | 16-NH | | 8.22, d (8.2) |
| Haorn-1 | 18 | 170.5, C | |
| | 19 | 55.6, CH | 4.27, m |
| | 20 | 29.8, $CH_2$ | 1.77, 1.58, m |
| | 21 | 20.1, $CH_2$ | 1.64, m |
| | 22 | 50.4, $CH_2$ | 3.07, t (5.3) |
| | 19-NH | | 8.07, d (8.2) |
| Ser-3 | 23 | 171.4, C | |
| | 24 | 55.6, CH | 4.33, m |
| | 25 | 62.4, $CH_2$ | 3.56, m |
| | 24-NH | | 8.00, d (8.2) |
| Haorn-2 | 26 | 165.2, C | |
| | 27 | 50.1, CH | 4.29, m |
| | 28 | 28.0, $CH_2$ | 1.86, 1.63, m |
| | 29 | 20.8, $CH_2$ | 1.88, m |
| | 30 | 51.8, $CH_2$ | 3.45, br |
| | 27-NH | | 8.23, d (8.2) |

Compound **5**, obtained as amorphous oil, has a molecular formula of $C_{14}H_{18}N_2O_7$ deduced by the HRESIMS. Analysis of the 1D and TOCSY NMR data. Analyses of COSY and TOCSY data reveled two amino acid spin systems consistent with a threonine derived moiety, one serine derived moiety. The coupling constants of H-2 (d, $J$ = 8.1 Hz), H-3 (t, $J$ = 7.6 Hz), H-4 (t, $J$ = 7.6 Hz) and H-5 (d, $J$ = 8.1 Hz) and COSY correlation of H-2/H-3/H-4/H-5, together with the HMBC correlations from H-5 to C-1, C-6 and C-7 indicated the presence of salicylic acid substructure. The amino acid sequence was assigned using a combination of HMBC, NOESY and TOCSY correlations, and MS/MS experiments. The first conjunction between salicylic acid substructure and threonine were supported by HMBC correlations from NH-9 to C-7 and C-9, as well as TOCSY correlation of NH-9/H-9/H-10/H-11. Furthermore, HMBC correlation from NH-13 to C-8 and C-13 and the TOCSY correlation of NH-13/H-13/H-14 indicated that serine-1 was attached to C-8 through N-13. Hence, the planar structure of **5** was determined. To deal with the absolute configuration of **5**, Marfey's method was performed by using L-FDAA to react with **5** and the result showed the configuration was 9*R*, 10*R*, 13*R*.

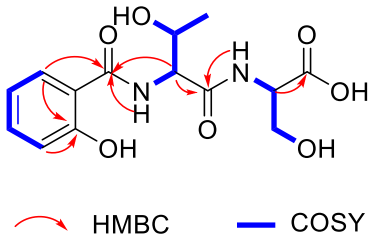


**Table 5**. $^{1}$H NMR (500 MHz) and $^{13}$C NMR (125 MHz) of **5** in DMSO-$d_6$ ($\delta$ in ppm, $J$ in Hz)

| **5** Unit | Position | $\delta_C$ type | $\delta_H$ ($J$ in Hz) |
|---|---|---|---|
| SA | 1 | 158.2, C | |
| | 2 | 130.3, CH | 7.93, d (8.3) |
| | 3 | 133.7, CH | 7.37, t (7.6) |
| | 4 | 119.6, CH | 6.92, t (7.8) |
| | 5 | 117.4, CH | 6.93, d (7.9) |
| | 6 | 117.8, C | |
| | 7 | 166.9, C | |
| Thr-1 | 8 | 170.3, C | |
| | 9 | 58.9, CH | 4.53, m |
| | 10 | 67.2, CH | 4.10, m |
| | 11 | 20.3, $CH_3$ | 1.07, d (6.4) |
| | 9-NH | | 8.79, d (8.2) |
| Ser-1 | 12 | 172.3, C | |
| | 13 | 55.1, CH | 4.30, m |
| | 14 | 61.8, $CH_2$ | 3.71, m |
| | 13-NH | | 8.06, d (8.2) |

Compound **6**, obtained as amorphous oil, has a molecular formula of $C_{14}H_{18}N_2O_7$ deduced by the HRESIMS. It was same as **5** by comparing with the NMR data. The only difference was the conjunction between salicylic acid substructure and threonine moiety, which was that O-10 was attached to C-7 through an ester bond, supported by the key HMBC correlation from H-10 to C-7 and the hydrogen integral of $NH_2$-9 was **6**. Hence, the planar structure of **6** was determined and the absolute configuration was 9*R*, 10*R*, 13*R*.

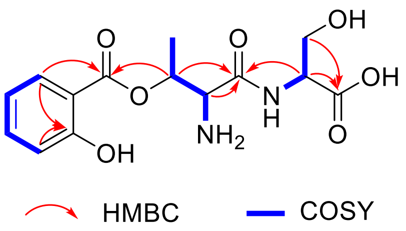


**Table 6**. [1]H NMR (400 MHz) and [13]C NMR (150 MHz) of **6** in DMSO-$d_6$ ($\delta$ in ppm, $J$ in Hz)

| **6** Unit | Position | $\delta_C$ type | $\delta_H$ ($J$ in Hz) |
|---|---|---|---|
| SA | 1 | 160.7, C | |
| | 2 | 131.7, CH | 7.96, d (8.3) |
| | 3 | 136.5, CH | 7.52, t (7.6) |
| | 4 | 119.7, CH | 6.93, t (7.8) |
| | 5 | 117.7, CH | 6.97, d (7.9) |
| | 6 | 113.4, C | |
| | 7 | 168.0, C | |
| Thr | 8 | 166.4, C | |
| | 9 | 55.6, CH | 4.25, br |
| | 10 | 70.5, CH | 5.42, m |
| | 11 | 16.8, $CH_3$ | 1.38, d (6.5) |
| | 9-NH | | 8.50, d (8.2) |
| Ser | 12 | 171.8, C | |
| | 13 | 55.2, CH | 4.35, dt (8.7, 4.5) |
| | 14 | 61.8, $CH_2$ | 3.70, dd (10.7, 4.8) |
| | 13-NH | | 8.90, d (8.2) |

Compound **7**, obtained as amorphous oil, has a molecular formula of $C_{14}H_{18}N_2O_7$ deduced by the HRESIMS. Analysis of NMR data indicated that **7** was same as with madurastatins B1. The only difference was the carboxyl group at C-8 is replaced by a serine, supported by the HMBC of NH-13 to C-8. Hence, the planar structure of **7** was determined. Considering the same biosynthesis pathway, the absolute configuration of **7** were deduced as 9*R*, 10*R*, 13*R*.

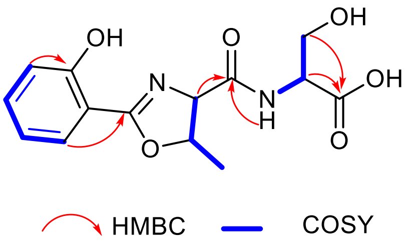


**Table 7**. [1]H NMR (400 MHz) and [13]C NMR (125 MHz) of **7** in DMSO-$d_6$ ($\delta$ in ppm, $J$ in Hz)

| **7** Unit | Position | $\delta_C$ type | $\delta_H$ ($J$ in Hz) |
|---|---|---|---|
| Cyc-SA | 1 | 160.7, C | |
| | 2 | 131.6, CH | 7.99, d (8.3) |
| | 3 | 136.5, CH | 7.55, t (7.6) |
| | 4 | 119.8, CH | 6.96, t (7.8) |
| | 5 | 117.1, CH | 7.01, d (7.9) |
| | 6 | 113.4, C | |
| | 7 | 168.0, C | |
| Cyc-Thr | 8 | 166.4, C | |
| | 9 | 55.6, CH | 4.26, br |
| | 10 | 70.5, CH | 5.45, m |
| | 11 | 16.8, $CH_3$ | 1.41, d (6.5) |
| Ser | 12 | 171.8, C | |
| | 13 | 55.3, CH | 4.34, m |
| | 14 | 61.8, $CH_2$ | 3.72, m |
| | 13-NH | | 9.00, d (8.2) |

Compound **8**, obtained as white powders, has a molecular formula of $C_{14}H_{18}N_2O_7$ deduced by the HRESIMS. Comparison of NMR data revealed that compounds 8 was similar with cittilins A. The only difference was the conjunction between 3-OH-Tyr$^2$ and 4-$CH_3$-Tyr$^3$ were through ester bond, supported by the unsaturation calculation and molecular formula based on high resolution calculation. Hence, the planar structure of **8** was determined. According to the literature, this type of compound was derived from ribosomal peptides, and its configuration is a natural L-amino acid. Therefore, the absolute configuration of the compound is tentatively determined to be 2*S*, 11*S*, 12*S*, 17*S*, 26*S*.

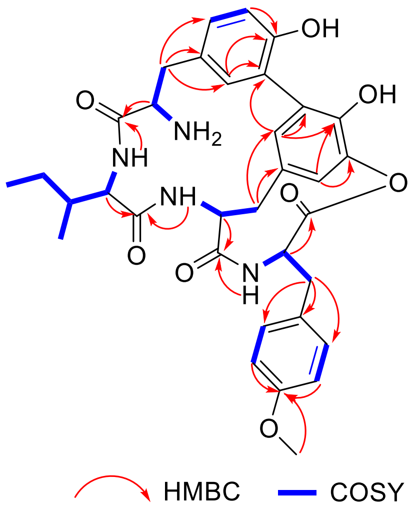

**Table 8**. $^{1}H$ NMR (400 MHz) and $^{13}C$ NMR (125 MHz) of **8** in DMSO-$d_6$ ($\delta$ in ppm, $J$ in Hz)

| **8** Unit | Position | $\delta_C$ type | $\delta_H$ ($J$ in Hz) |
|---|---|---|---|
| Tyr-1 | 1 | 167.8, C | |
| | 2 | 52.5, CH | 4.13, m |
| | 3 | 35.8, $CH_2$ | 3.24, d (14.0), 2.99 d (10.1) |
| | 4 | 124.9, C | |
| | 5 | 131.1, CH | 7.17, m |
| | 6 | 111.0, CH | 6.96, d (8.4) |
| | 7 | 156.5, C | |
| | 8 | 128.2, C | |
| | 9 | 135.9, CH | 6.71, s |
| | 2-$NH_2$ | | 7.89, s |
| Ile-1 | 10 | 171.0, C | |
| | 11 | 57.9, CH | 4.18, m |
| | 12 | 36.2, CH | 1.64, m |
| | 13 | 25.1, $CH_2$ | 1.46, m, 1.15, m |
| | 14 | 11.2, $CH_3$ | 0.83, m |
| | 15 | 15.9, $CH_3$ | 0.85, m |
| | 11-NH | | 8.64, d (7.6) |
| OH-Tyr-2 | 16 | 171.3, C | |
| | 17 | 58.2, CH | 3.48, m |
| | 18 | 40.1, $CH_2$ | 2.67, d (12.6), 2.41, d (13.2) |
| | 19 | 134.5, C | |
| | 20 | 119.0, CH | 5.52, s |
| | 21 | 152.7, C | |
| | 22 | 128.2, C | |
| | 23 | 144.5, C | |
| | 24 | 125.4, CH | 6.41, s |
| | 17-NH | | 8.72, d (9.4) |
| Me-Tyr-2 | 25 | 173.0, C | |
| | 26 | 51.1, CH | 4.32, m |
| | 27 | 35.4, $CH_2$ | 3.38, m, 3.13, m |
| | 28 | 132.8, C | |
| | 29 | 135.0, CH | 7.33, d (7.8) |
| | 30 | 124.2, CH | 6.80, d (6.6) |
| | 31 | 163.0, C | |
| | 32 | 126.6, CH | 7.55, d (6.4) |
| | 33 | 129.8, CH | 7.16, m |
| | 34 | 55.9, $CH_3$ | 3.78, s |
| | 26-NH | | 8.93, s |

#### Physicochemical Property

##### *Nocsiderin* A (1)

amorphous oil; UV (MeOH) $\lambda_{max}$ 220 (1.67), 280 (1.15) nm; $^{1}$H and $^{13}$C NMR data, Table S1; HRESIMS *m/z* 743.3201 $[M + H]^{+}$ (calcd for $C_{30}H_{47}N_{8}O_{14}$, 743.3212).

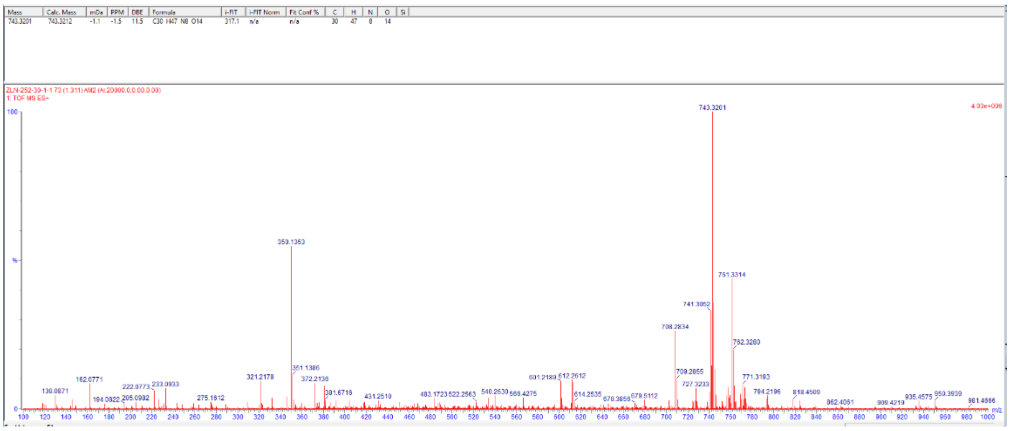


##### *Nocsiderin* B (2)

amorphous oil; UV (MeOH) $\lambda_{max}$ 220 (1.65), 280 (1.15) nm; $^{1}$H and $^{13}$C NMR data, Table S2; HRESIMS *m/z* 743.3200 $[M + H]^{+}$ (calcd for $C_{14}H_{23}O_{3}$, 743.3206).

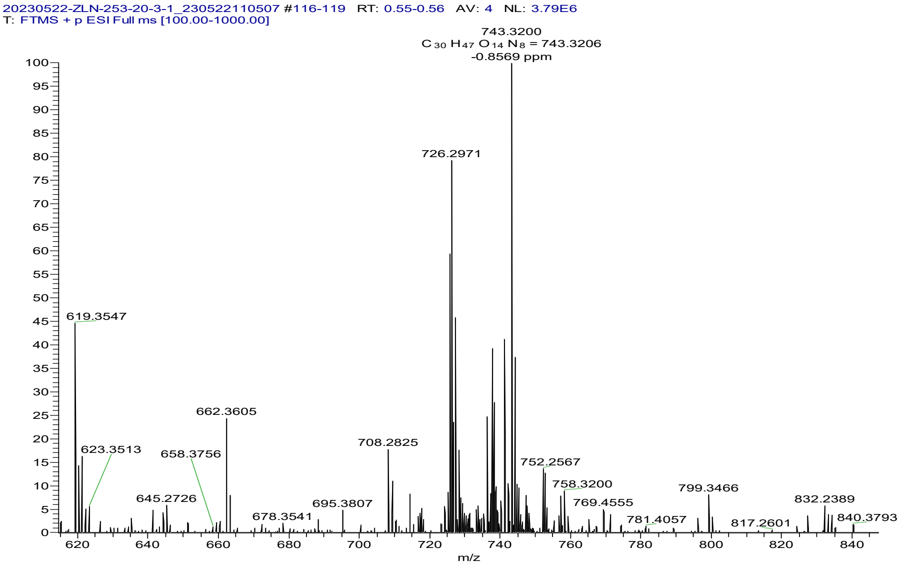


##### *Nocsiderin* C (3)

amorphous oil; UV (MeOH) $\lambda_{max}$ 220 (1.67), 280 (1.15) nm; $^1$H and $^{13}$C NMR data, Table S3; HRESIMS *m/z* 771.3171 [M + H]$^+$ (calcd for $C_{17}H_{21}O_3$, 771.3161).

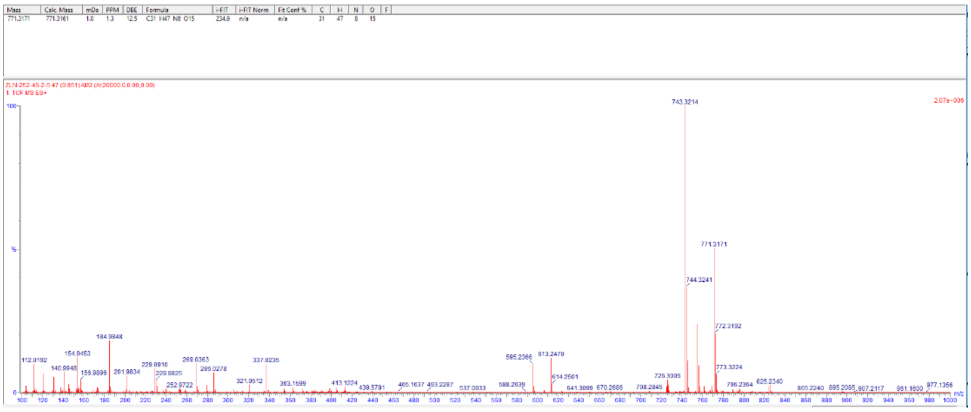

#### *Nocsiderin* D (4)

amorphous oil; UV (MeOH) $\lambda_{max}$ 220 (1.66), 280 (1.05) nm; $^1$H and $^{13}$C NMR data, Table S4; HRESIMS *m/z* 743.3209 [M + H]$^+$ (calcd for $C_{17}H_{21}O_3$, 743.3212).

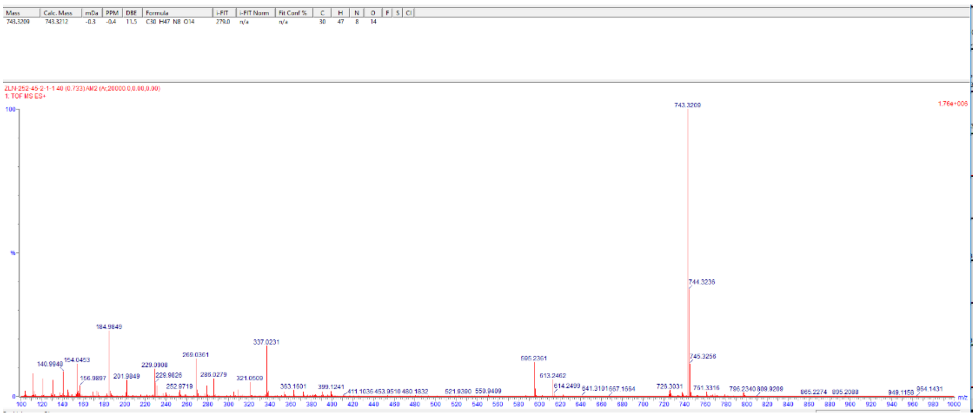

#### *Nocsiderin* E (5)

amorphous oil; UV (MeOH) $\lambda_{max}$ 220 (1.47), 280 (1.15) nm; $^1$H and $^{13}$C NMR data, Table S5; HRESIMS *m/z* 327.1199 [M + H]$^+$ (calcd for $C_{17}H_{21}O_3$, 327.1192).

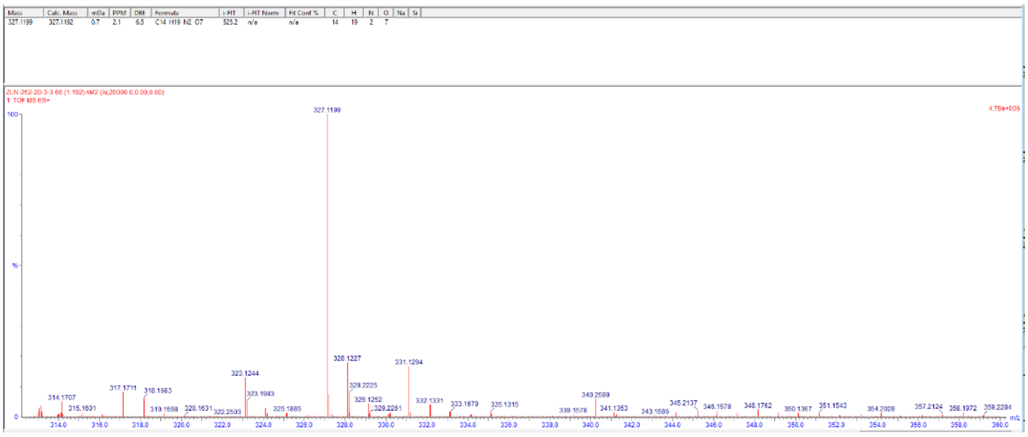


#### *Nocsiderin* F (6)

amorphous oil; UV (MeOH) $\lambda_{max}$ 220 (1.45), 280 (1.05) nm; $^{1}$H and $^{13}$C NMR data, Table S6; HRESIMS *m/z* 349.1013 $[M + Na]^+$ (calcd for $C_{17}H_{21}O_3$, 349.1012).

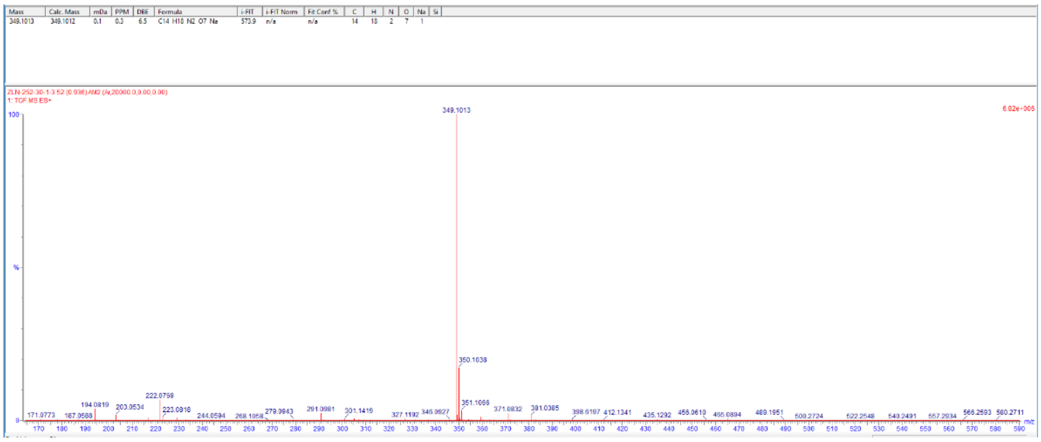


#### *Nocsiderin* G (7)

amorphous oil; UV (MeOH) $\lambda_{max}$ 220 (1.47), 280 (1.15) nm; $^{1}$H and $^{13}$C NMR data, Table S7; HRESIMS *m/z* 309.1093 $[M + H]^+$ (calcd for $C_{17}H_{21}O_3$, 309.1087).

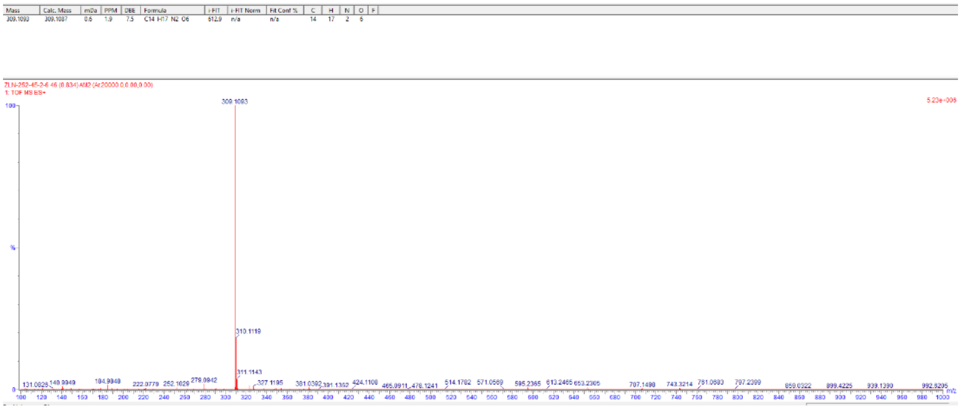


#### *Nocsiderin* G (8)

amorphous oil; UV (MeOH) $\lambda_{max}$ 220 (1.77), 280 (1.35) nm; $^{1}$H and $^{13}$C NMR data, Table S8; HRESIMS *m/z* 631.2771 $[M + H]^+$ (calcd for $C_{17}H_{21}O_3$, 631.2768).

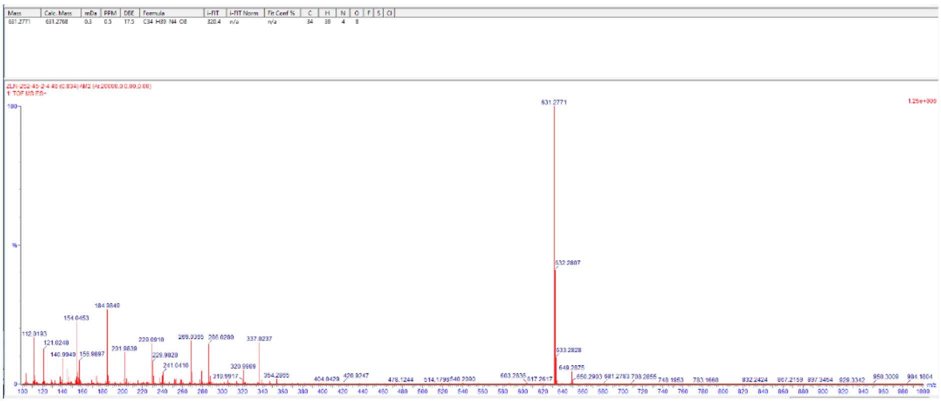


#### Marfey's results

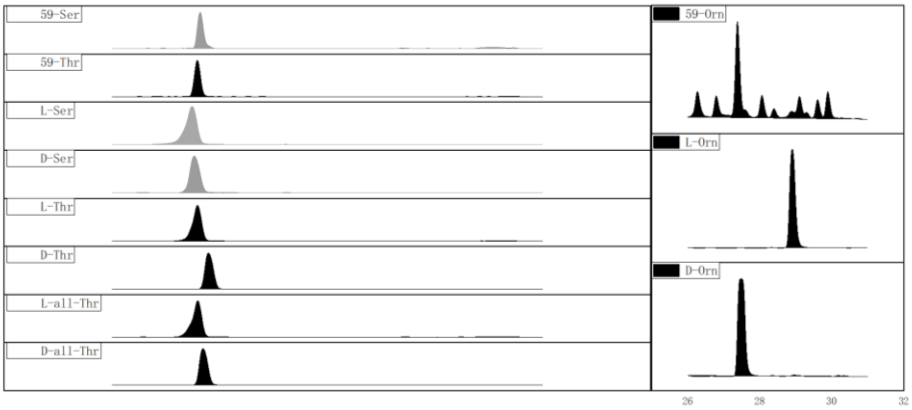

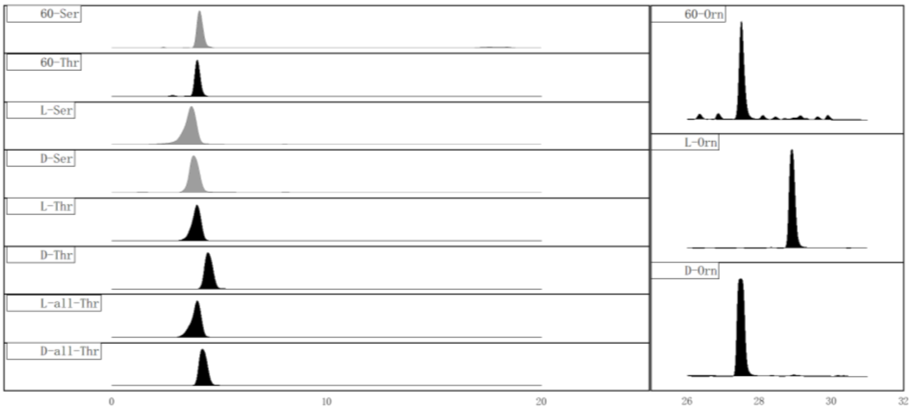
60-Ser
60-Thr
L-Ser
D-Ser
L-Thr
D-Thr
L-all-Thr
D-all-Thr
60-Orn
L-Orn
D-Orn
0
10
20
26
28
30
32

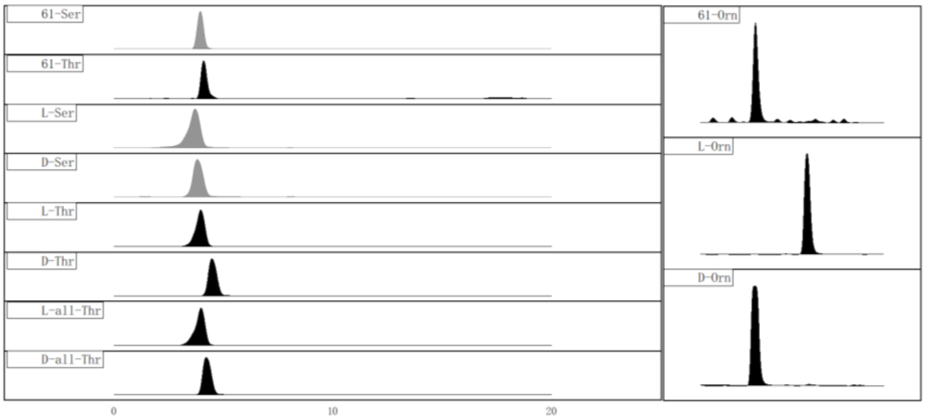
61-Ser
61-Thr
L-Ser
D-Ser
L-Thr
D-Thr
L-all-Thr
D-all-Thr
61-Orn
L-Orn
D-Orn
0
10
20

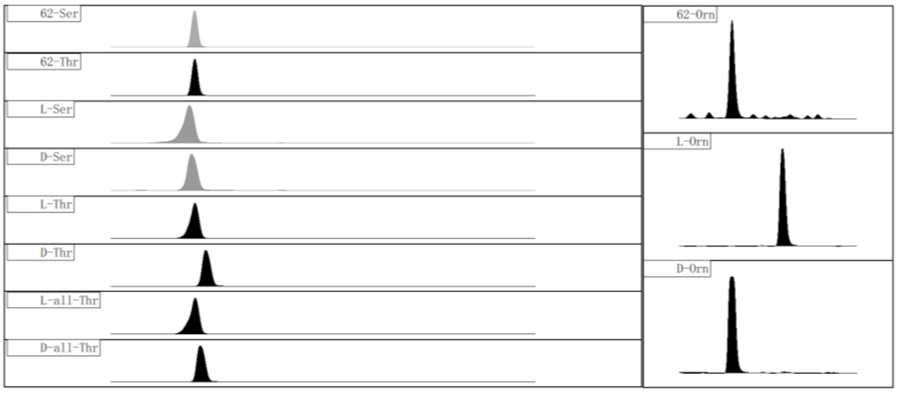
62-Ser
62-Thr
L-Ser
D-Ser
L-Thr
D-Thr
L-all-Thr
D-all-Thr
62-Orn
L-Orn
D-Orn

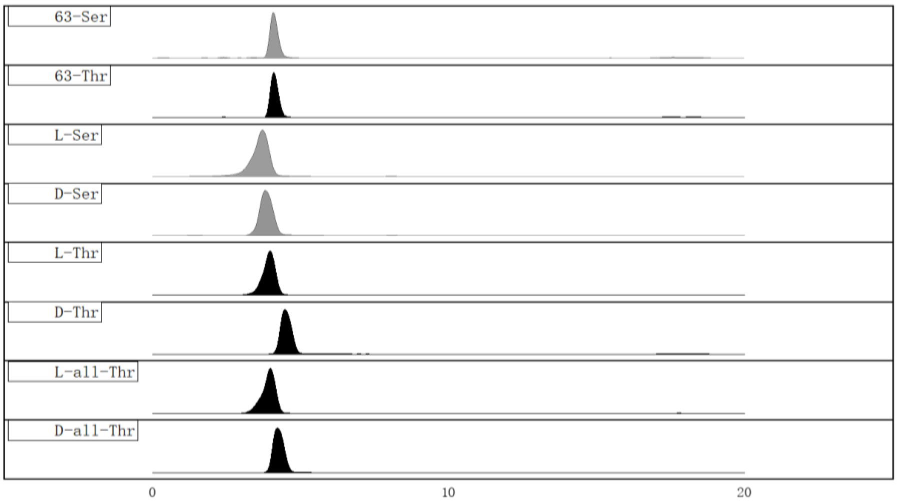
63-Ser
63-Thr
L-Ser
D-Ser
L-Thr
D-Thr
L-all-Thr
D-all-Thr
0
10
20

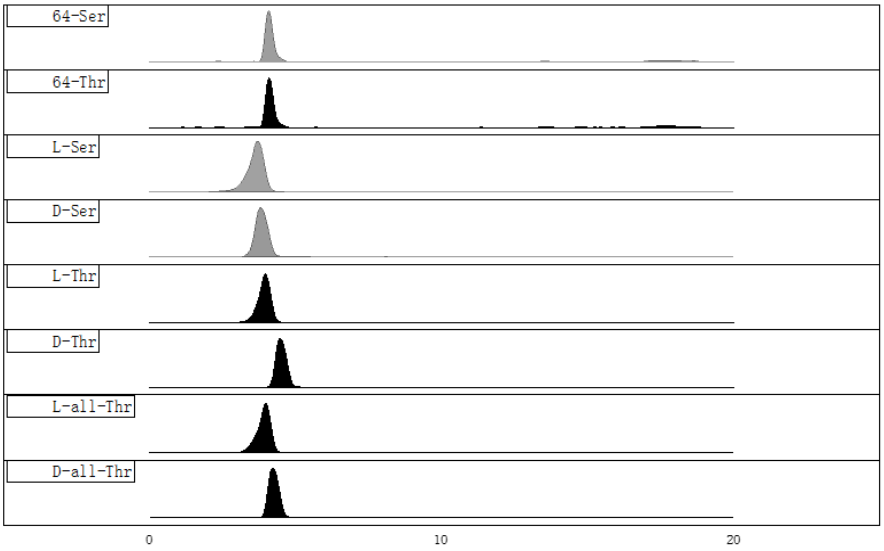
64-Ser
64-Thr
L-Ser
D-Ser
L-Thr
D-Thr
L-all-Thr
D-all-Thr
0
10
20

### NMR Spectrum Data

**Figure 1**. $^{1}$H NMR (400 MHz) spectrum of **1** in DMSO-$d_6$ ($\delta$ in ppm, $J$ in Hz)

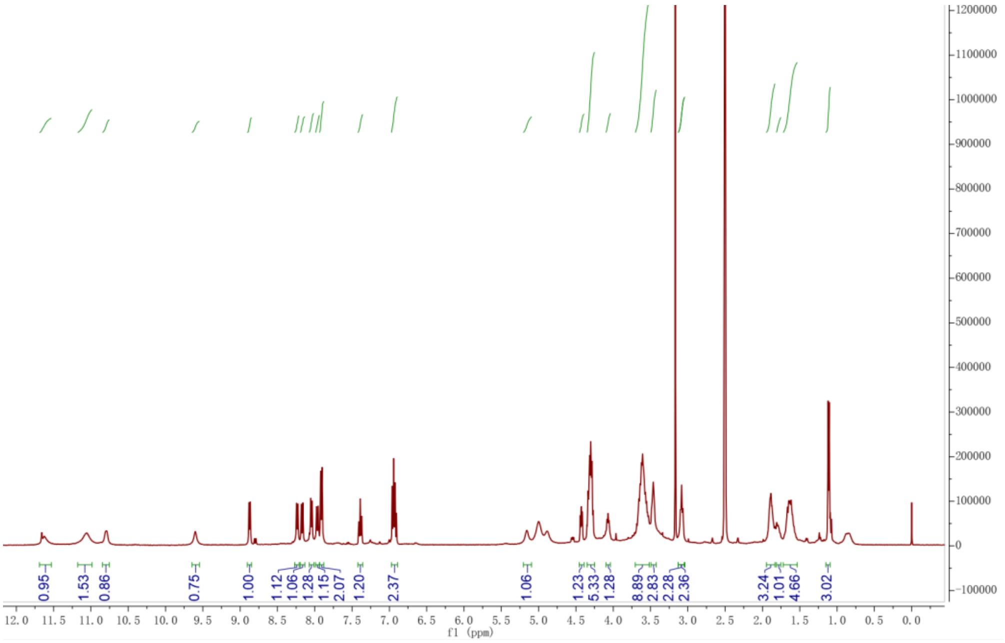


**Figure 2**. $^{13}$C NMR (150 MHz) spectrum of **1** in DMSO-$d_6$ ($\delta$ in ppm, $J$ in Hz)

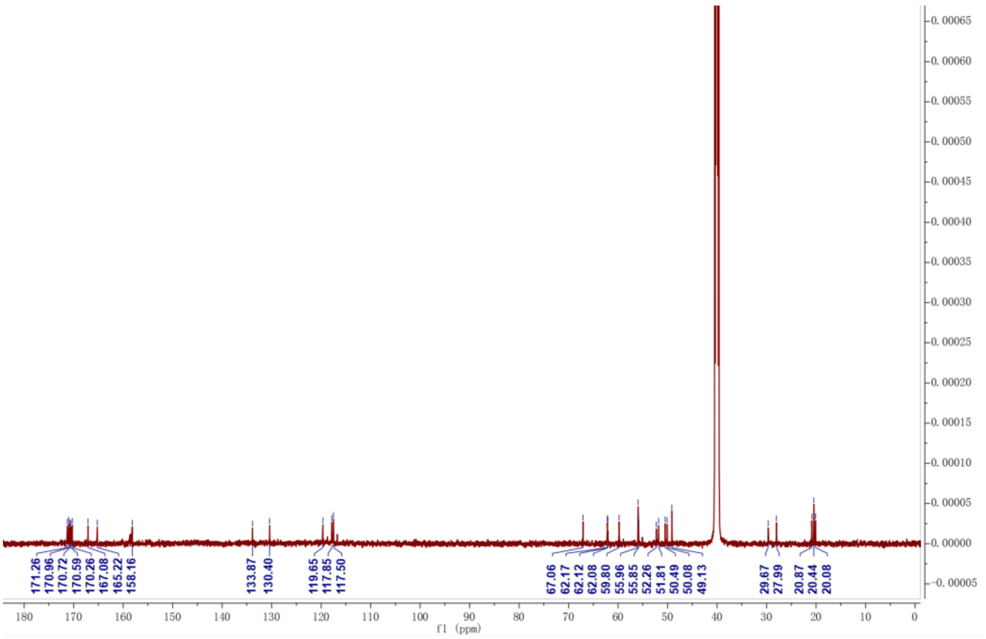


**Figure 3**. HSQC spectrum of **1** in DMSO-$d_6$ ($\delta$ in ppm, $J$ in Hz)

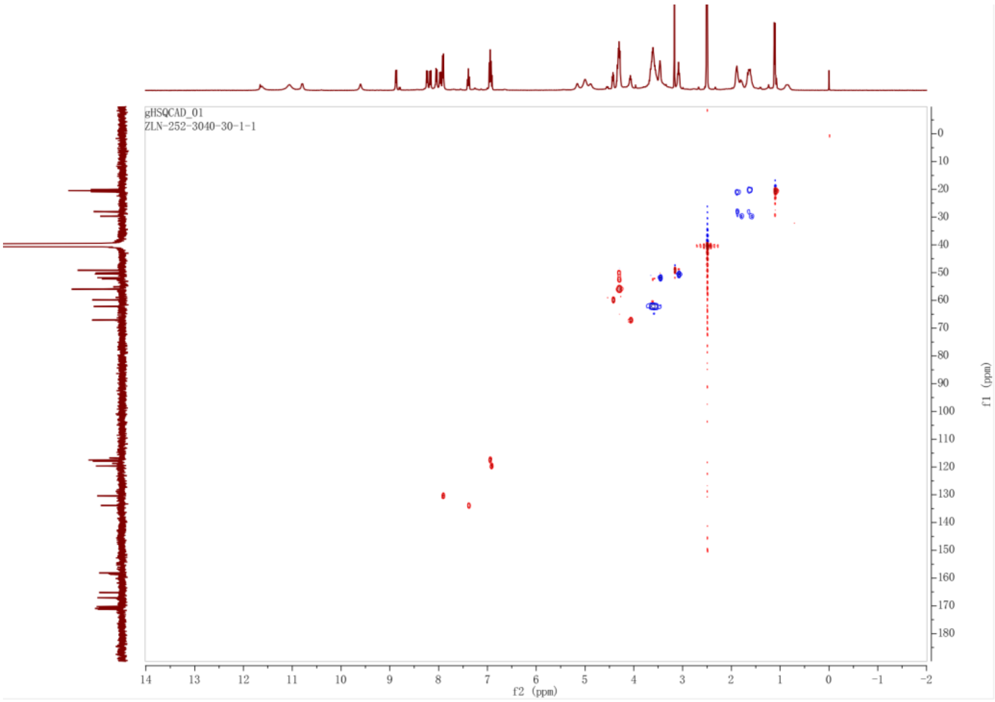


**Figure 4**. HMBC spectrum of **1** in DMSO-$d_6$ ($\delta$ in ppm, $J$ in Hz)

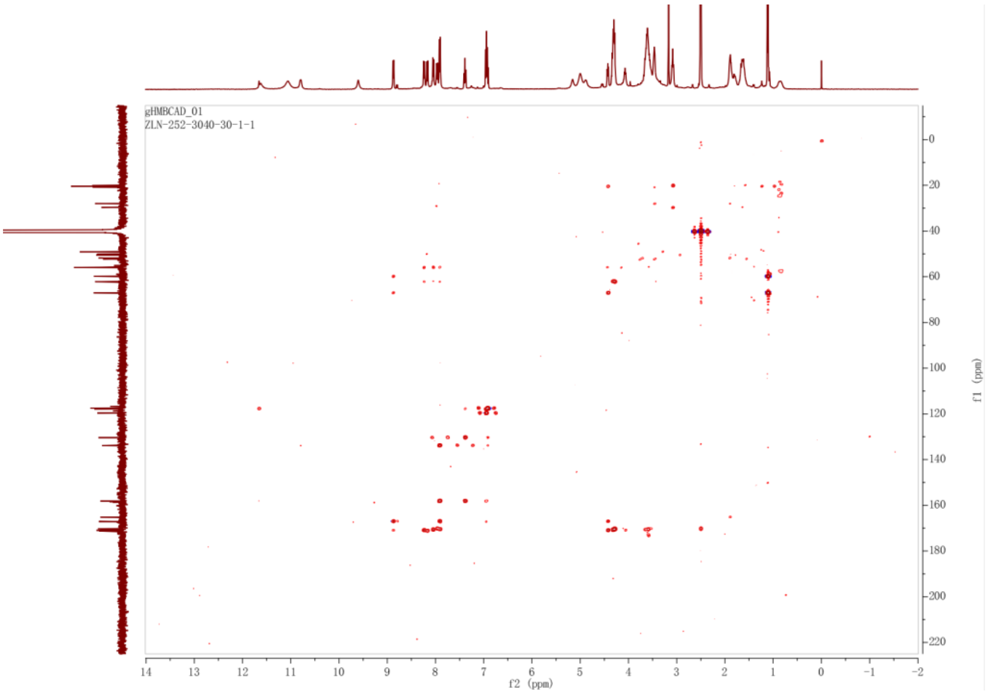


**Figure 5**. TOCSY spectrum of **1** in DMSO-$d_6$ ($\delta$ in ppm, $J$ in Hz)

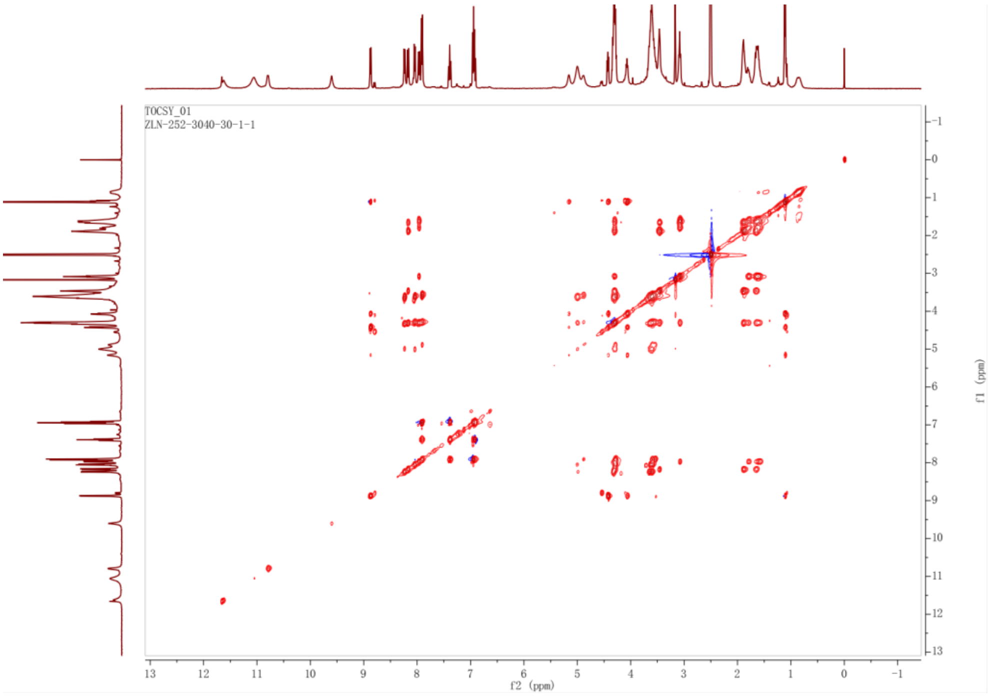


**Figure 5**. NOESY spectrum of **1** in DMSO-$d_6$ ($\delta$ in ppm, $J$ in Hz)

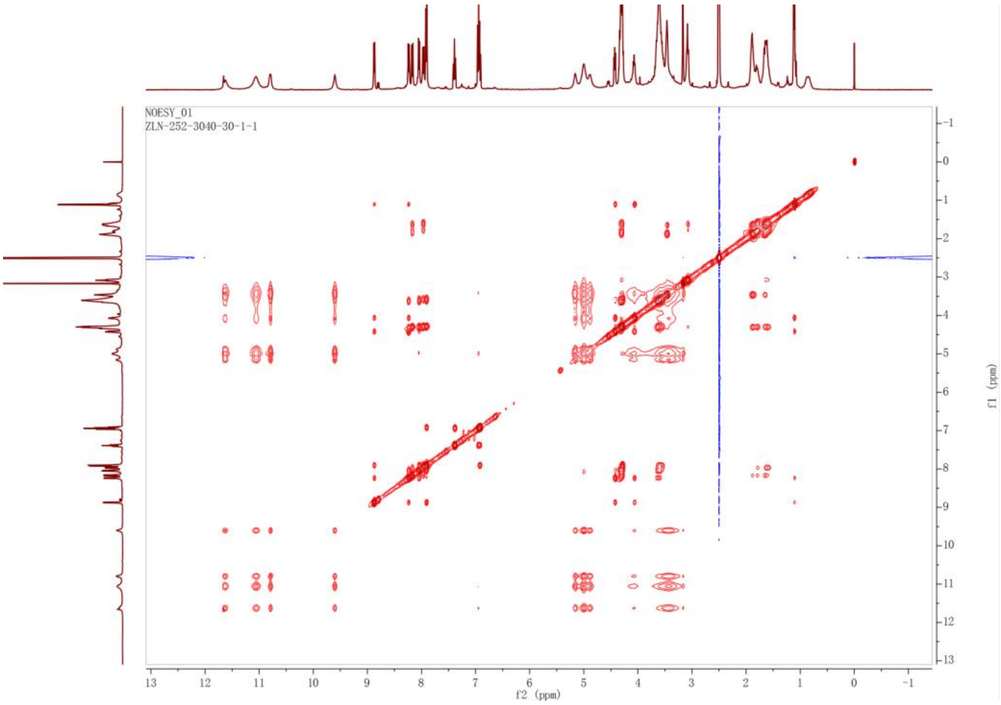


**Figure 6**. $^1$H NMR (400 MHz) spectrum of **2** in DMSO-$d_6$ ($\delta$ in ppm, $J$ in Hz)

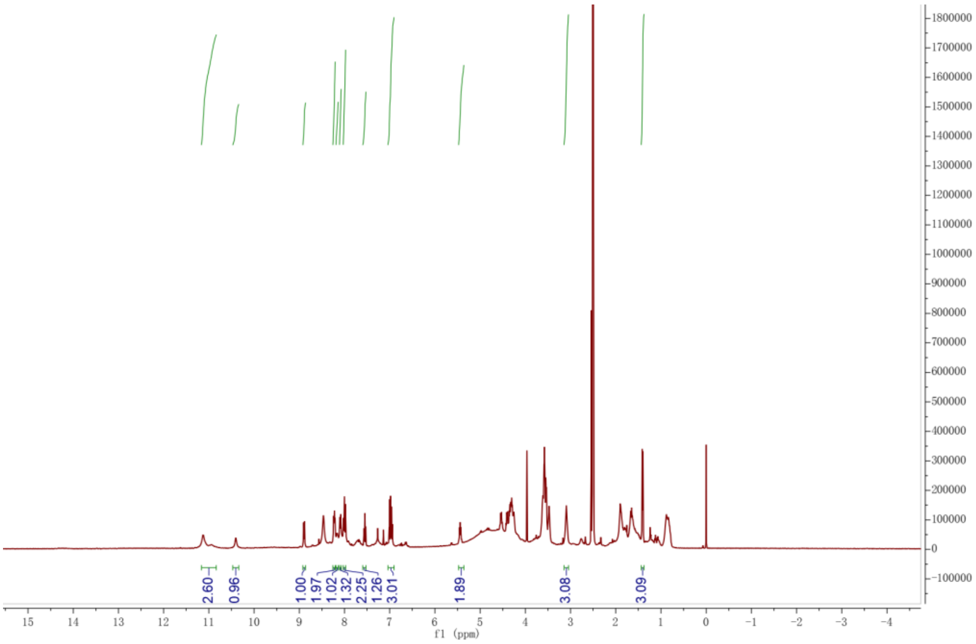


**Figure 8**. $^{13}$C NMR (125 MHz) spectrum of **2** in DMSO-$d_6$ ($\delta$ in ppm, $J$ in Hz)

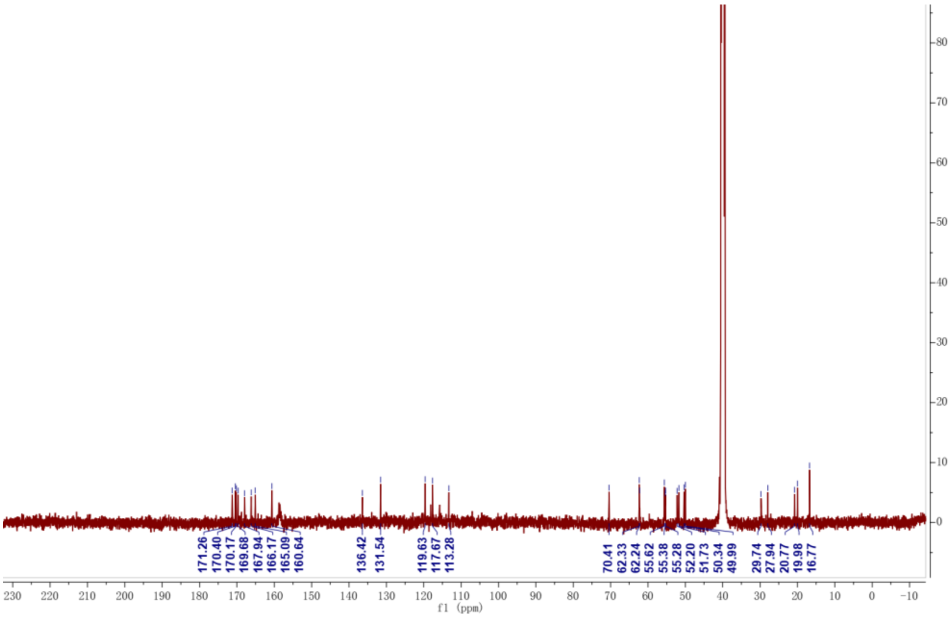


**Figure 9**. HSQC spectrum of **2** in DMSO-$d_6$ ($\delta$ in ppm, $J$ in Hz)

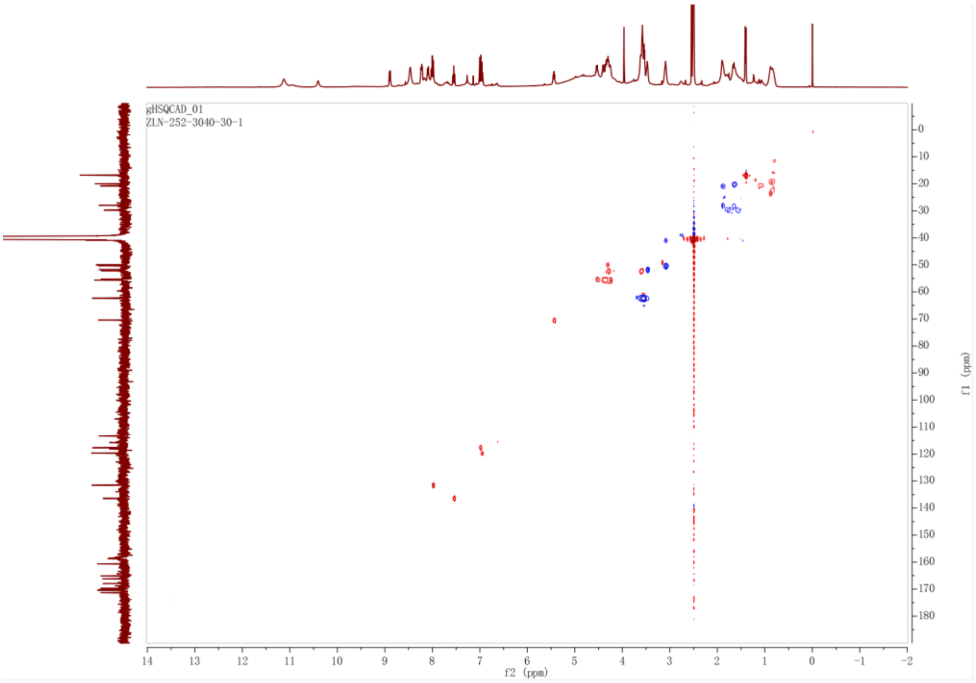


**Figure 10**. HMBC spectrum of **2** in DMSO-$d_6$ ($\delta$ in ppm, $J$ in Hz)

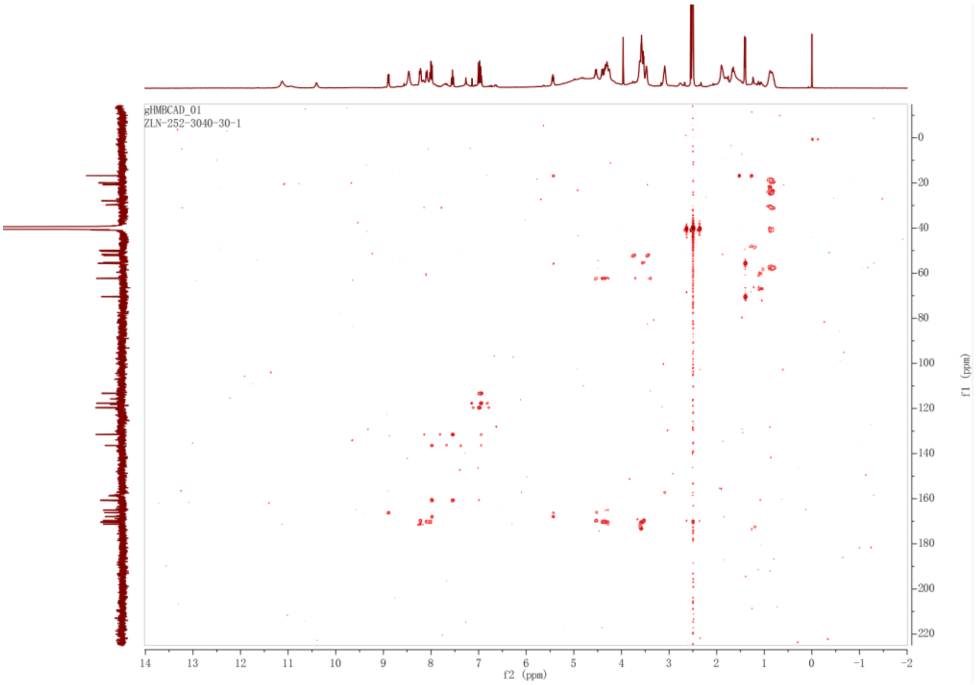


**Figure 11**. TOCSY spectrum of **2** in DMSO-$d_6$ ($\delta$ in ppm, $J$ in Hz)

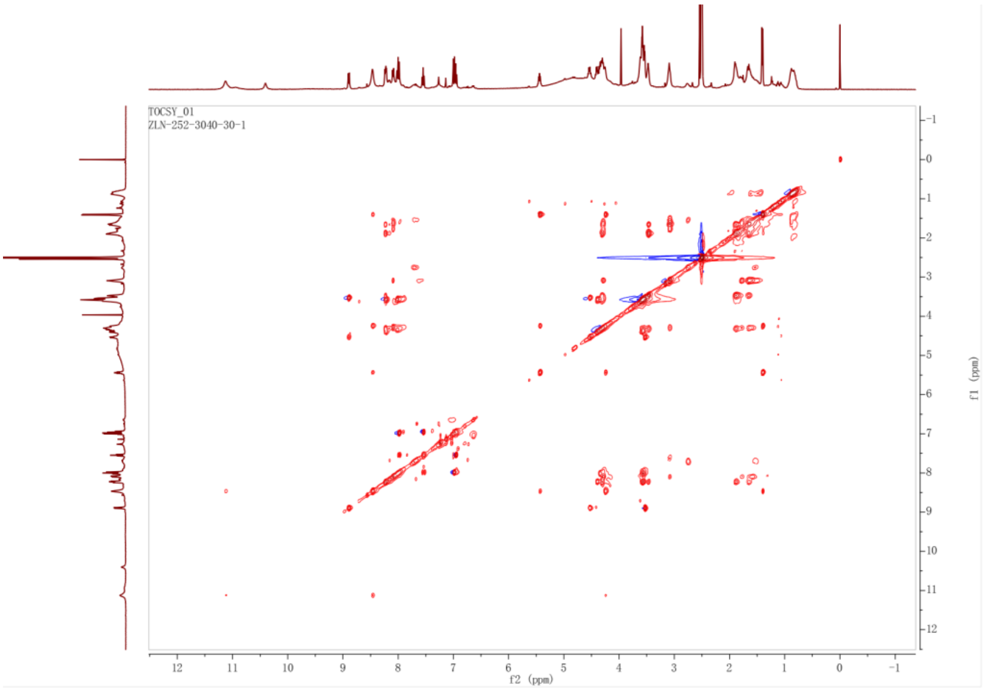


**Figure 12**. NOESY spectrum of **2** in DMSO-$d_6$ ($\delta$ in ppm, $J$ in Hz)

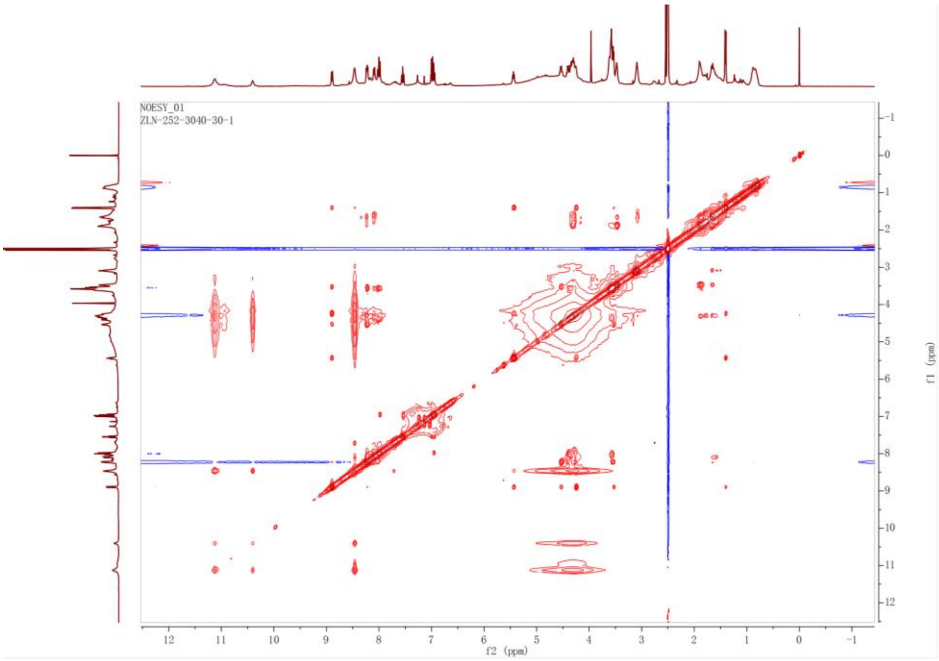


**Figure 13**. $^1$H NMR (400 MHz) spectrum of **3** in DMSO-$d_6$ ($\delta$ in ppm, $J$ in Hz)

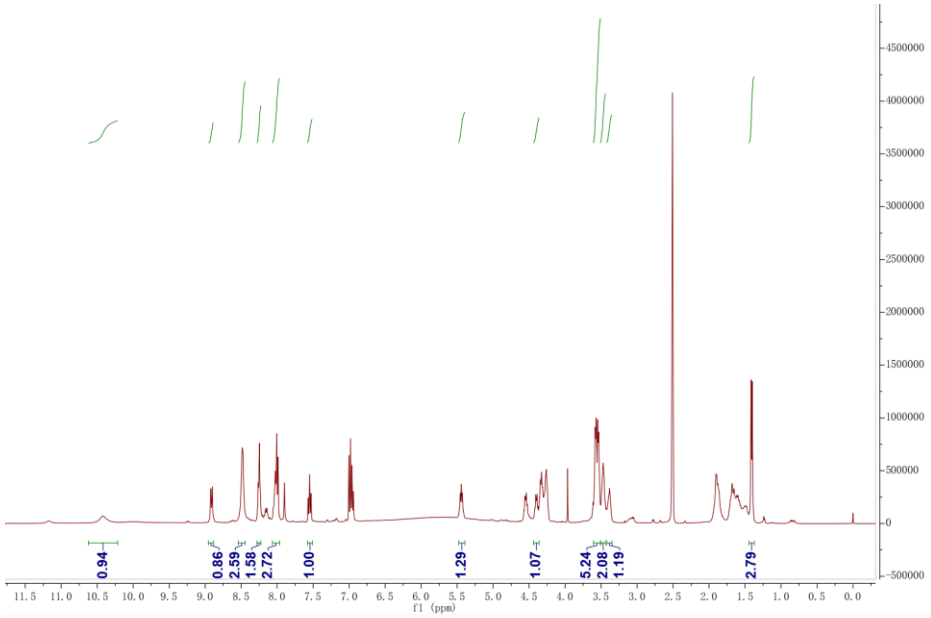


**Figure 14**. $^{13}$C NMR (125 MHz) spectrum of **3** in DMSO-$d_6$ ($\delta$ in ppm, $J$ in Hz)

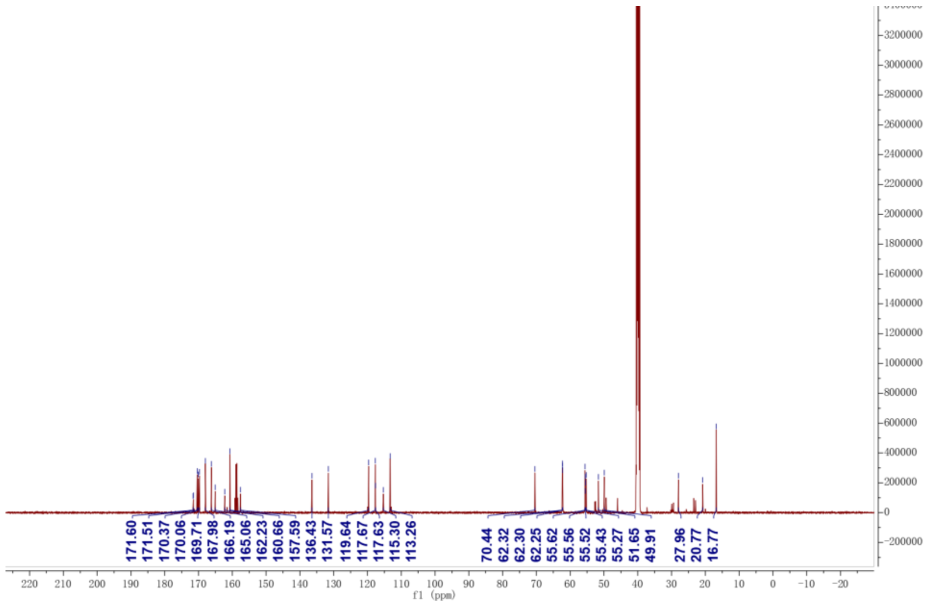


**Figure 15**. HSQC spectrum of **3** in DMSO-$d_6$ ($\delta$ in ppm, $J$ in Hz)

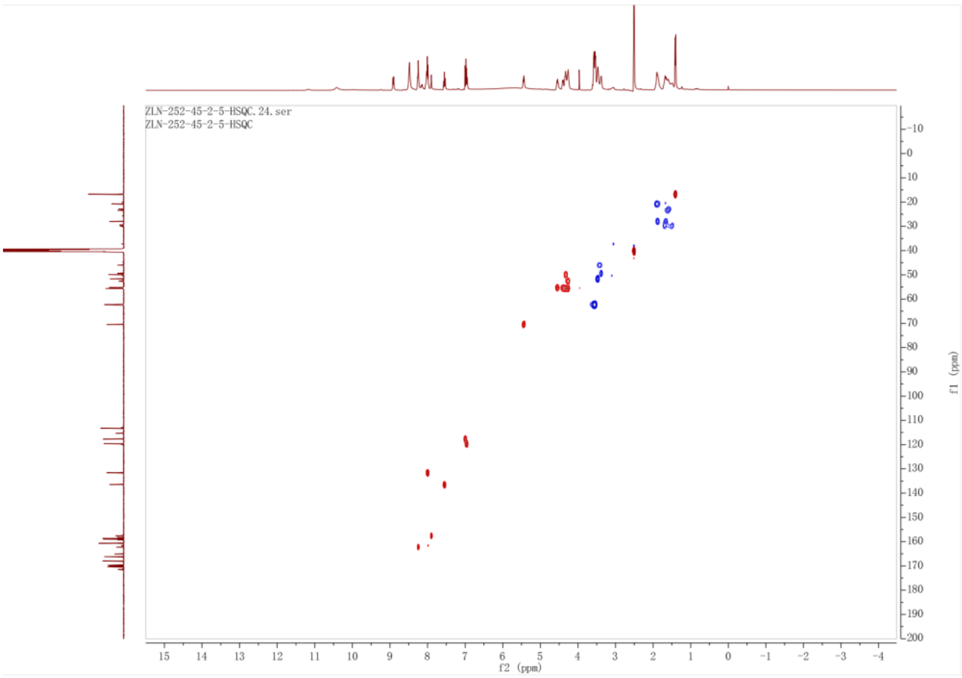


**Figure 16**. HMBC spectrum of **3** in DMSO-$d_6$ ($\delta$ in ppm, $J$ in Hz)

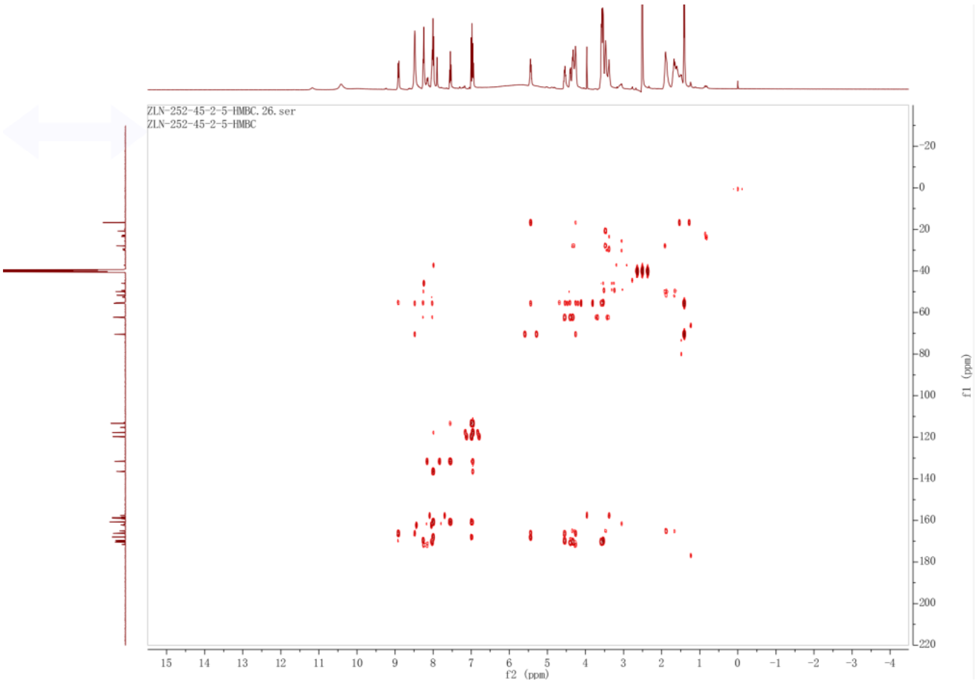


**Figure 17**. TOCSY spectrum of **3** in DMSO-$d_6$ ($\delta$ in ppm, $J$ in Hz)

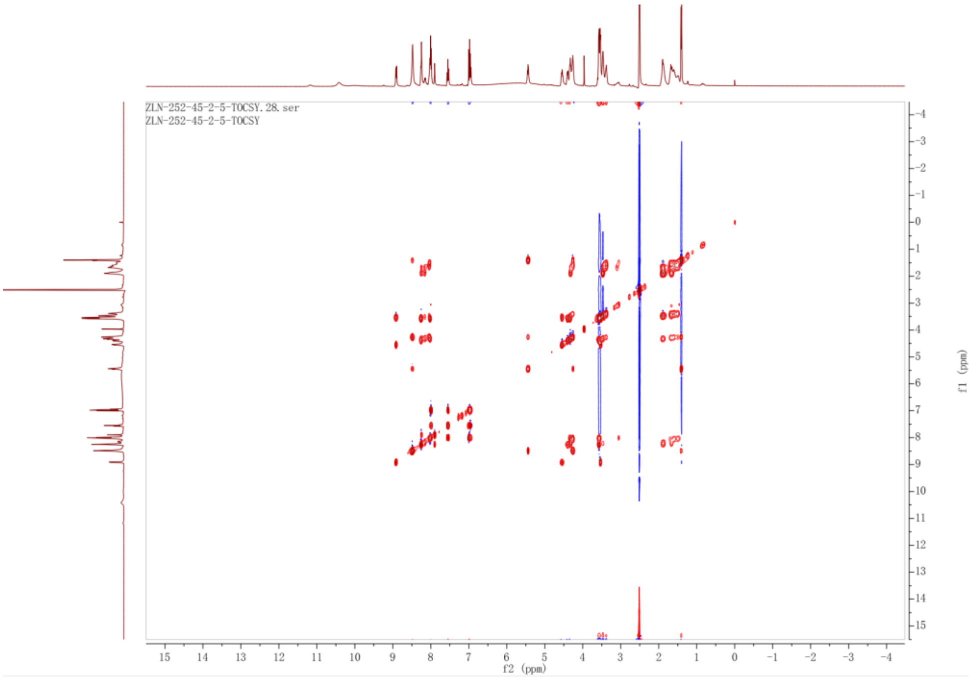


**Figure 18**. NOESY spectrum of **3** in DMSO-$d_6$ ($\delta$ in ppm, $J$ in Hz)

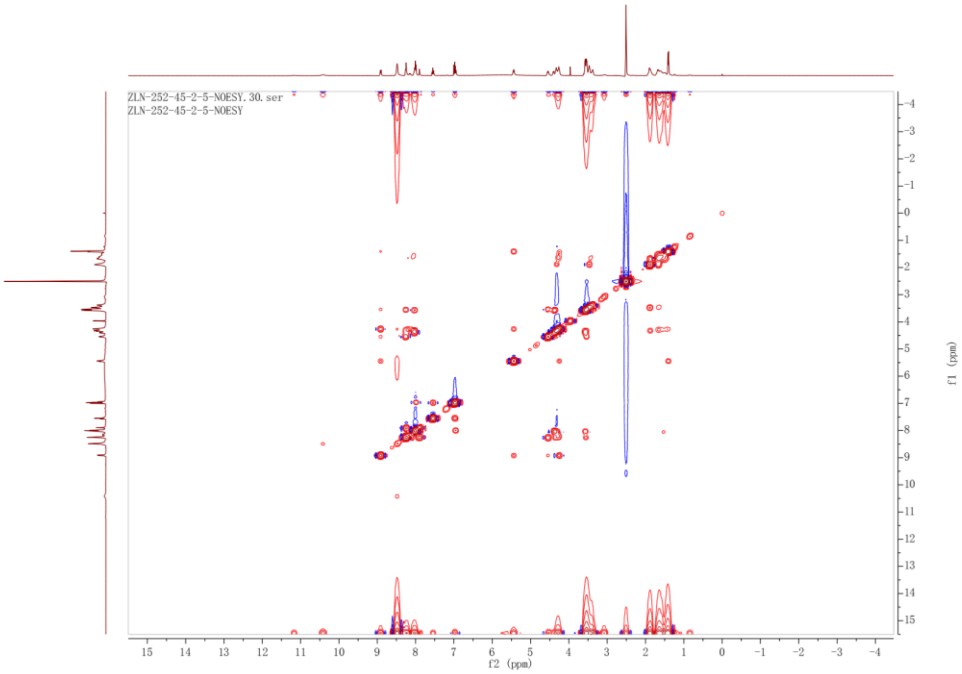


**Figure 19**. $^{1}$H NMR (500 MHz) spectrum of **4** in DMSO-$d_6$ ($\delta$ in ppm, $J$ in Hz)

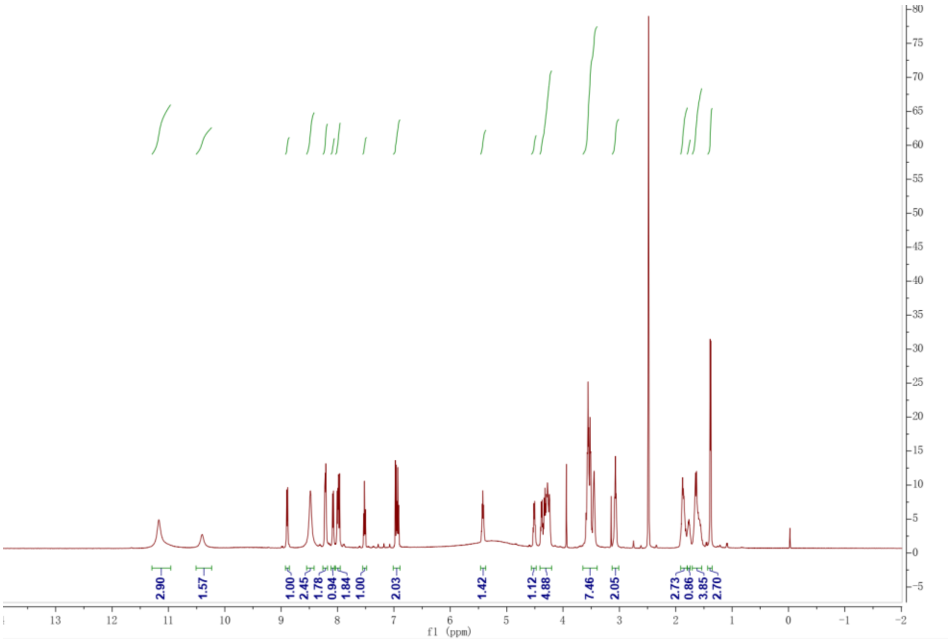

**Figure 20**. $^{13}$C NMR (150 MHz) spectrum of **4** in DMSO-$d_6$ ($\delta$ in ppm, $J$ in Hz)

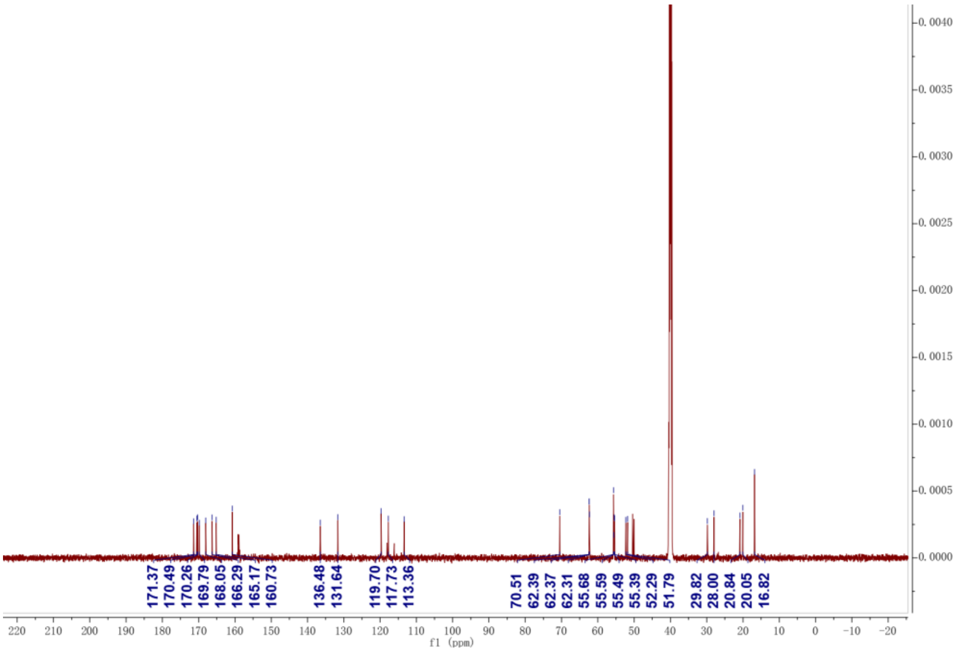

**Figure 21**. HSQC spectrum of **4** in DMSO-$d_6$ ($\delta$ in ppm, $J$ in Hz)

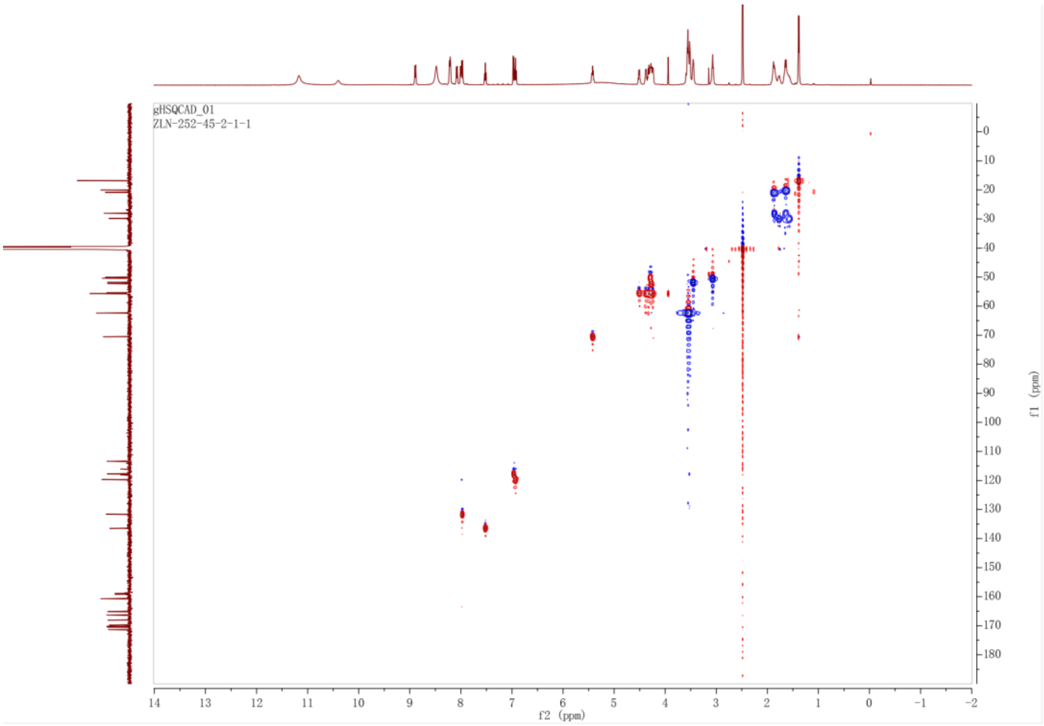


**Figure 22**. HMBC spectrum of **4** in DMSO-$d_6$ ($\delta$ in ppm, $J$ in Hz)

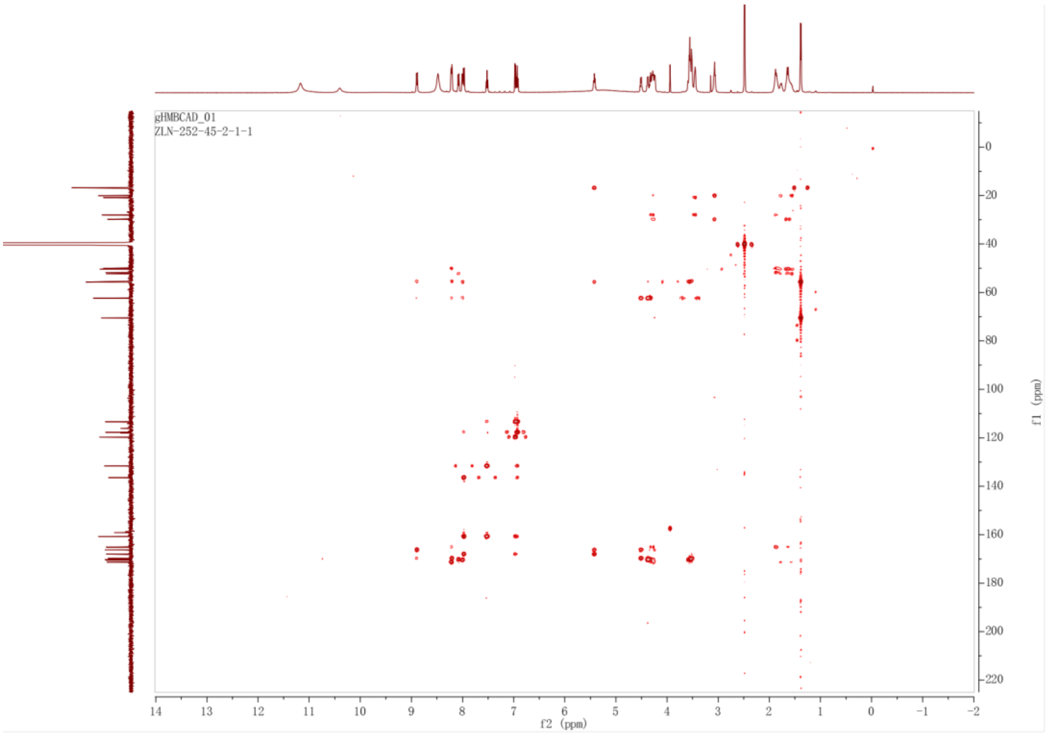


**Figure 23**. TOCSY spectrum of **4** in DMSO-$d_6$ ($\delta$ in ppm, $J$ in Hz)

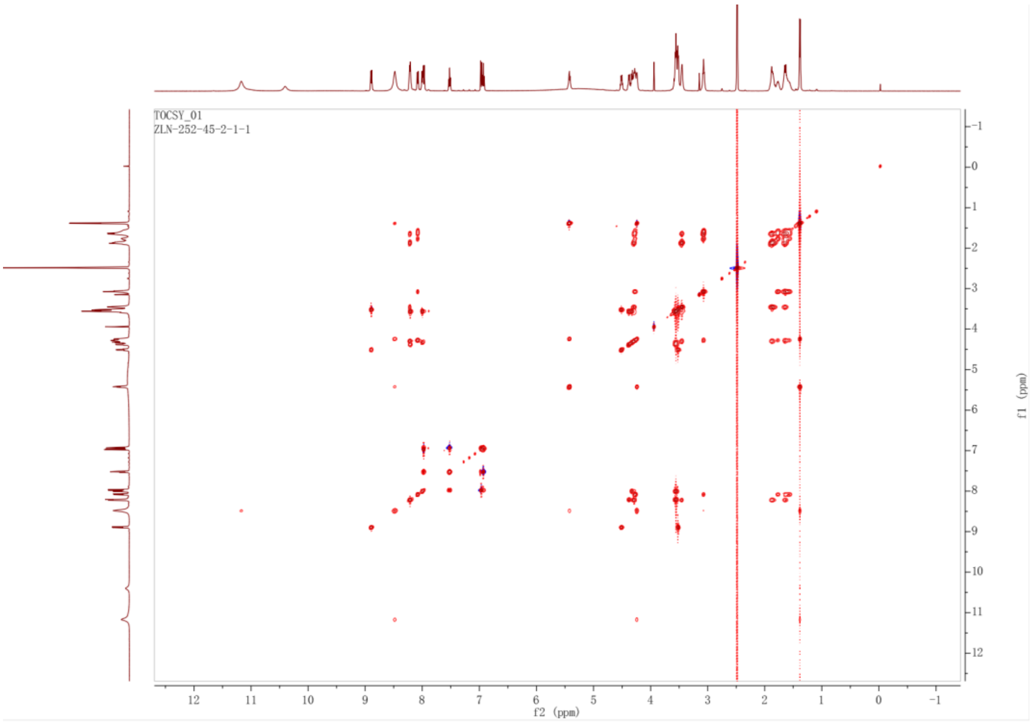


**Figure 24**. NOESY spectrum of **4** in DMSO-$d_6$ ($\delta$ in ppm, $J$ in Hz)

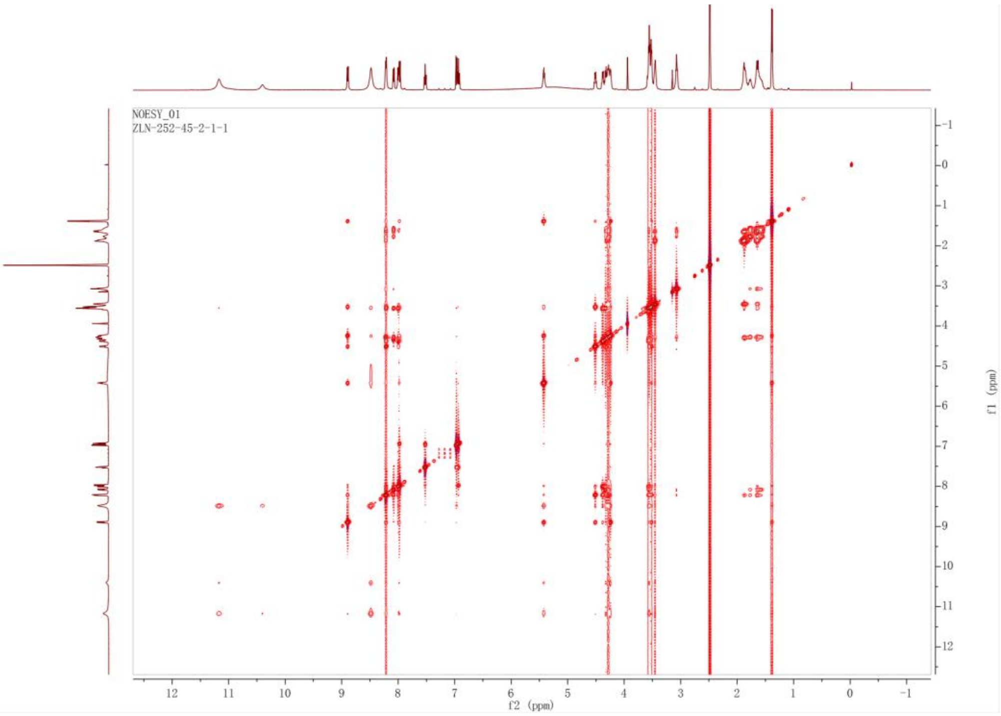


**Figure 25**. $^1$H NMR (500 MHz) spectrum of **5** in DMSO-$d_6$ ($\delta$ in ppm, $J$ in Hz)

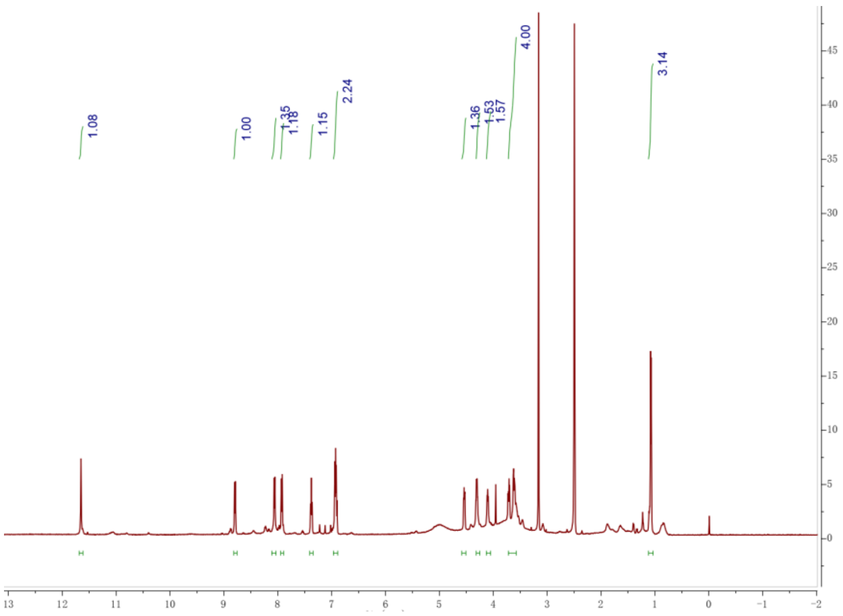

**Figure 26**. $^{13}$C NMR (125 MHz) spectrum of **5** in DMSO-$d_6$ ($\delta$ in ppm, $J$ in Hz)

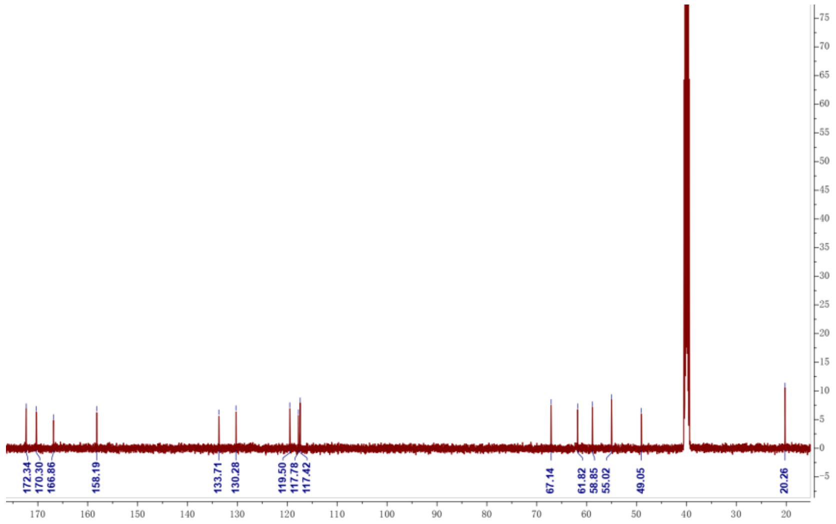


**Figure 27**. HSQC spectrum of **5** in DMSO-$d_6$ ($\delta$ in ppm, $J$ in Hz)

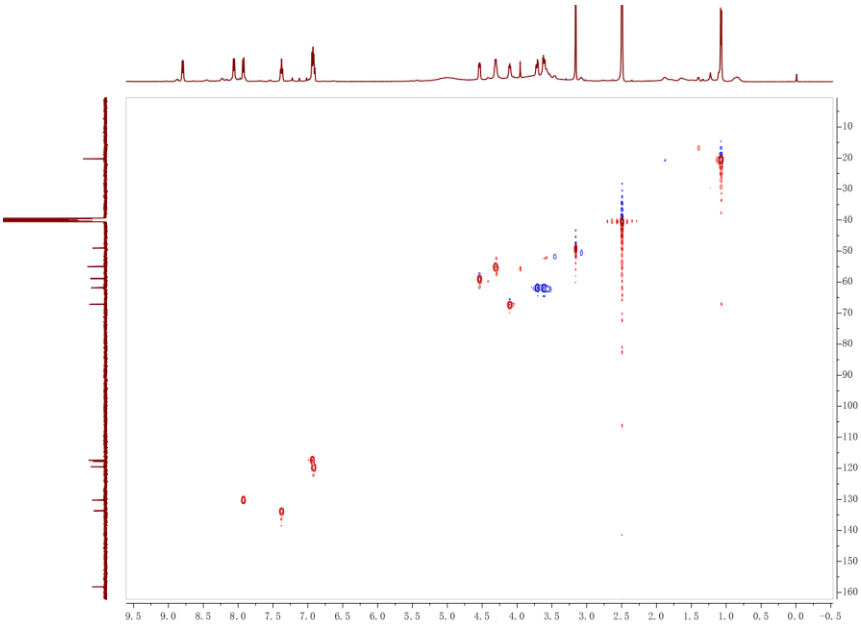


**Figure 28**. HMBC spectrum of **5** in DMSO-$d_6$ ($\delta$ in ppm, $J$ in Hz)

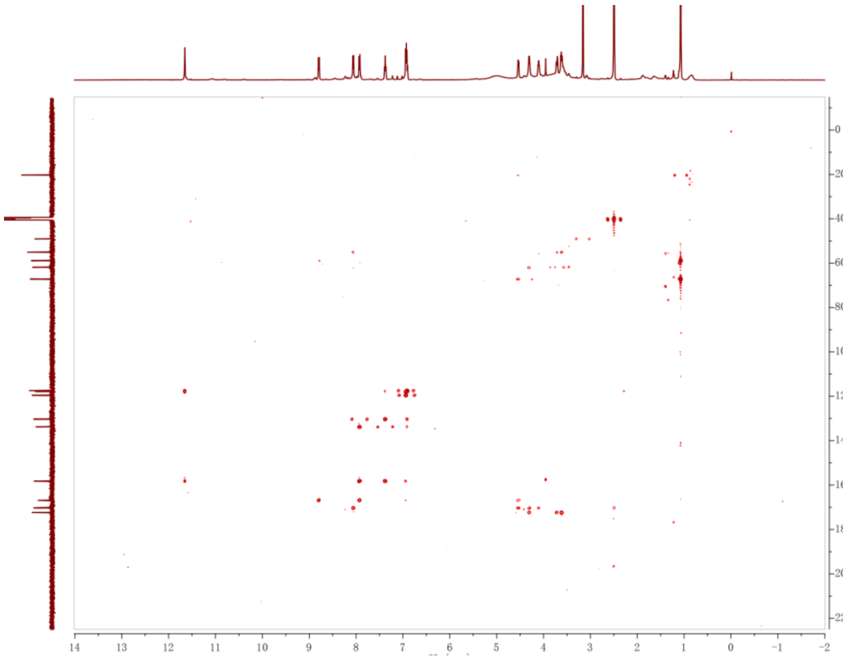


**Figure 29**. $^1$H NMR (400 MHz) spectrum of **6** in DMSO-$d_6$ ($\delta$ in ppm, $J$ in Hz)

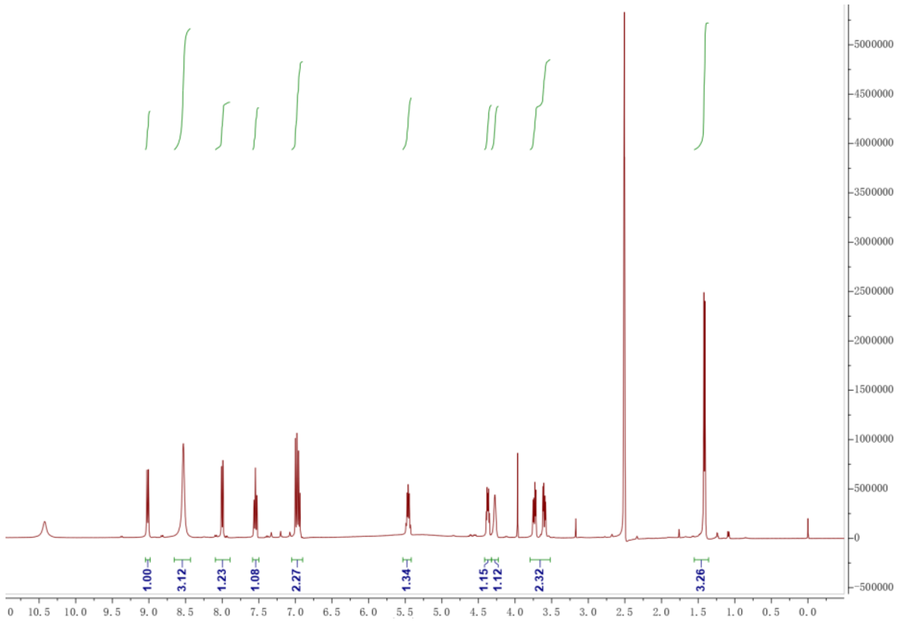


**Figure 30**. $^{13}$C NMR (150 MHz) spectrum of **6** in DMSO-$d_6$ ($\delta$ in ppm, $J$ in Hz)

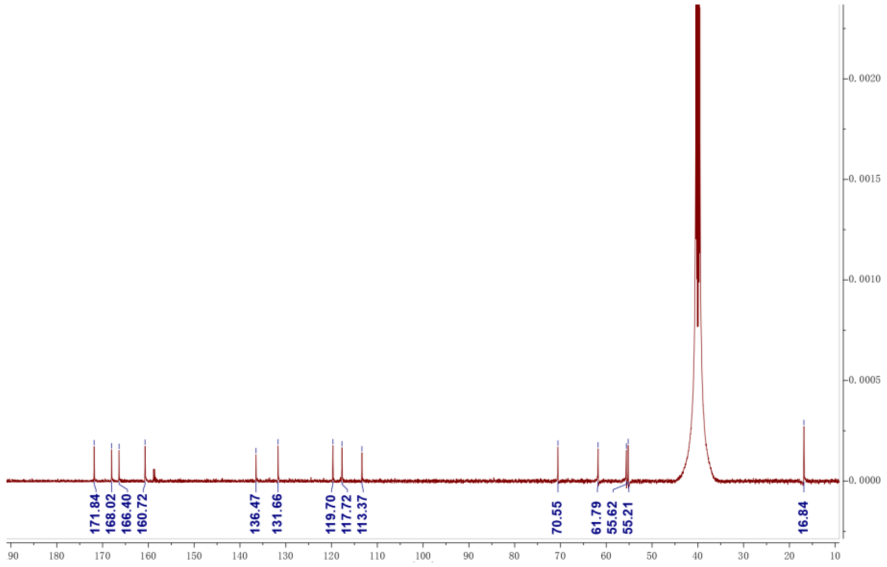


**Figure 31**. HSQC spectrum of **6** in DMSO-$d_6$ ($\delta$ in ppm, $J$ in Hz)

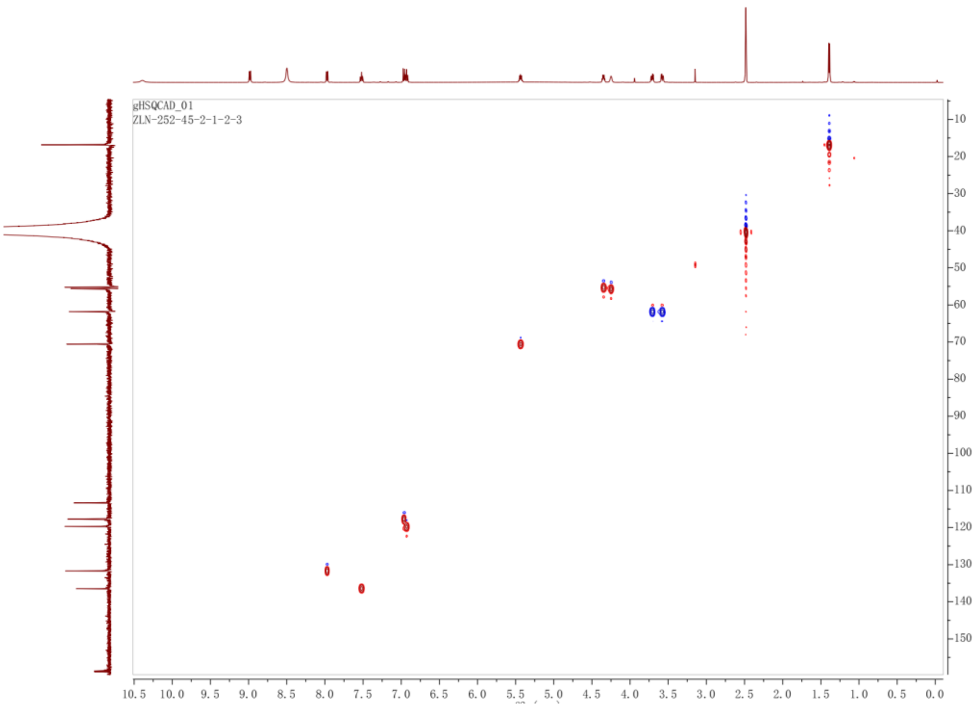


**Figure 32**. HMBC spectrum of **6** in DMSO-$d_6$ ($\delta$ in ppm, $J$ in Hz)

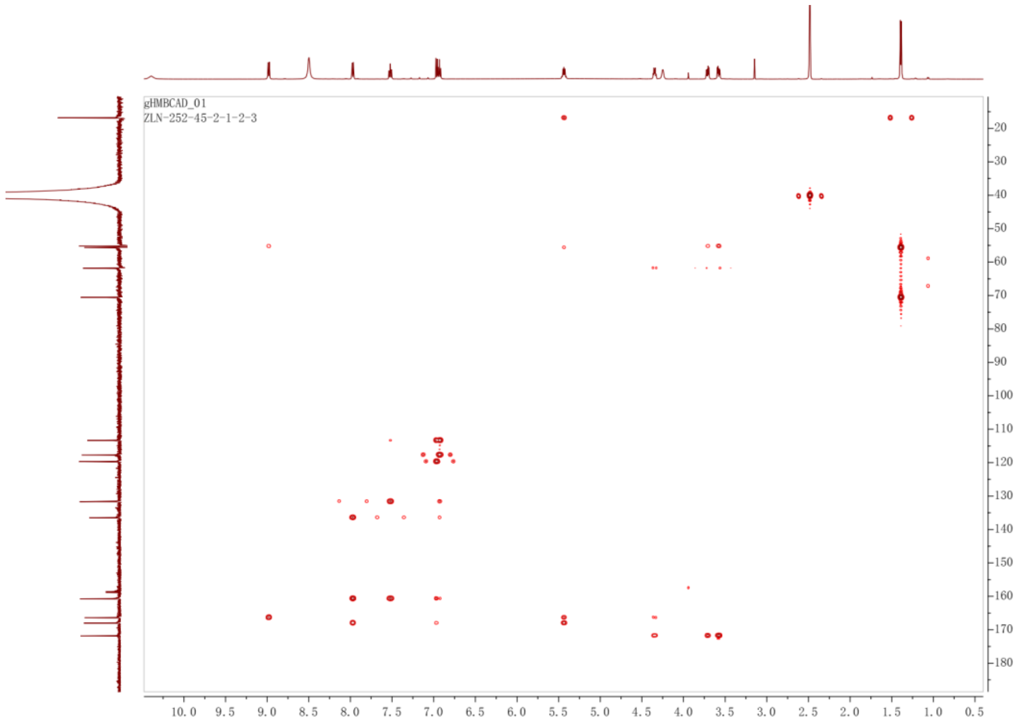


**Figure 33**. $^1$H NMR (400 MHz) spectrum of **7** in DMSO-$d_6$ ($\delta$ in ppm, $J$ in Hz)

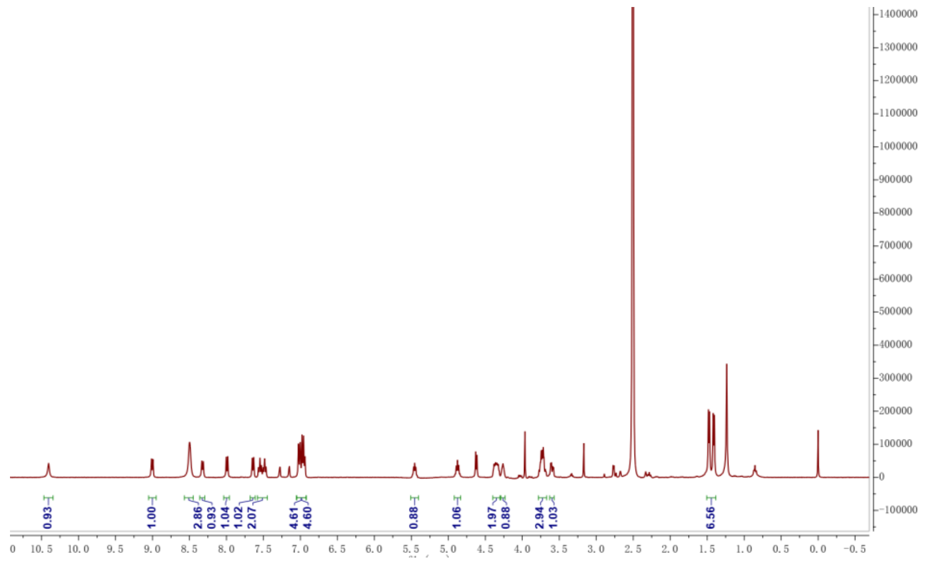


**Figure 34**. $^{13}$C NMR (125 MHz) spectrum of **7** in DMSO-$d_6$ ($\delta$ in ppm, $J$ in Hz)

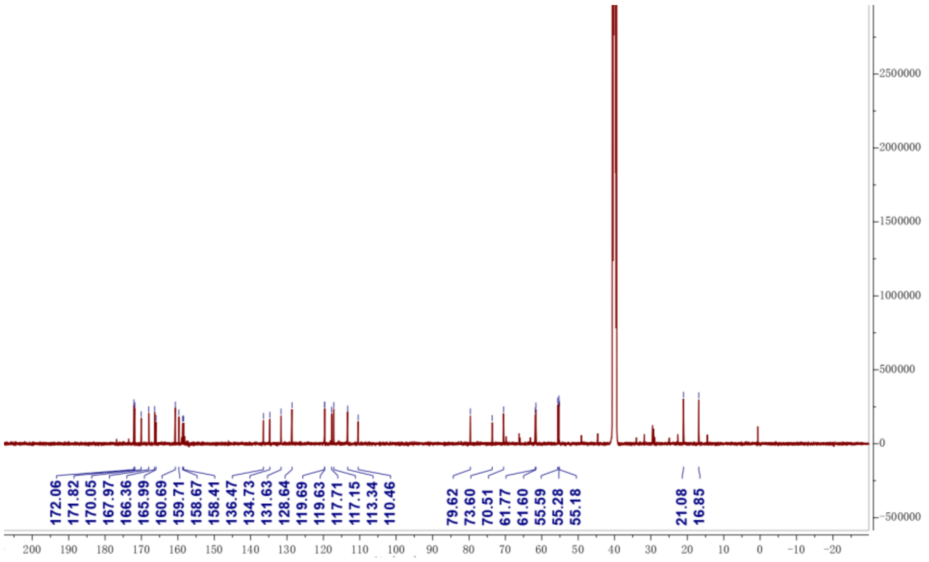


**Figure 35**. HSQC spectrum of **7** in DMSO-$d_6$ ($\delta$ in ppm, $J$ in Hz)

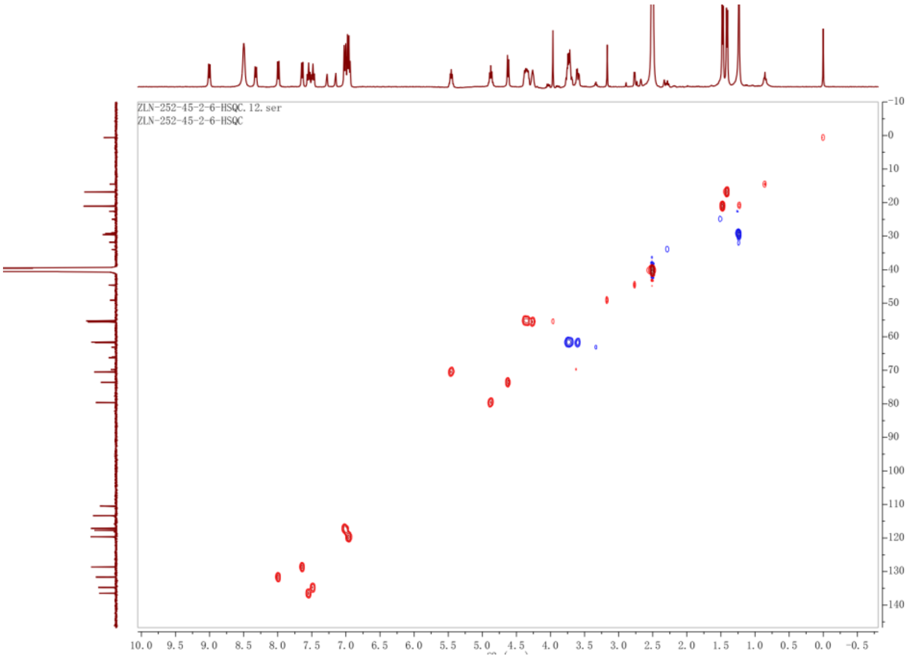


**Figure 36**. HMBC spectrum of **7** in DMSO-$d_6$ ($\delta$ in ppm, $J$ in Hz)

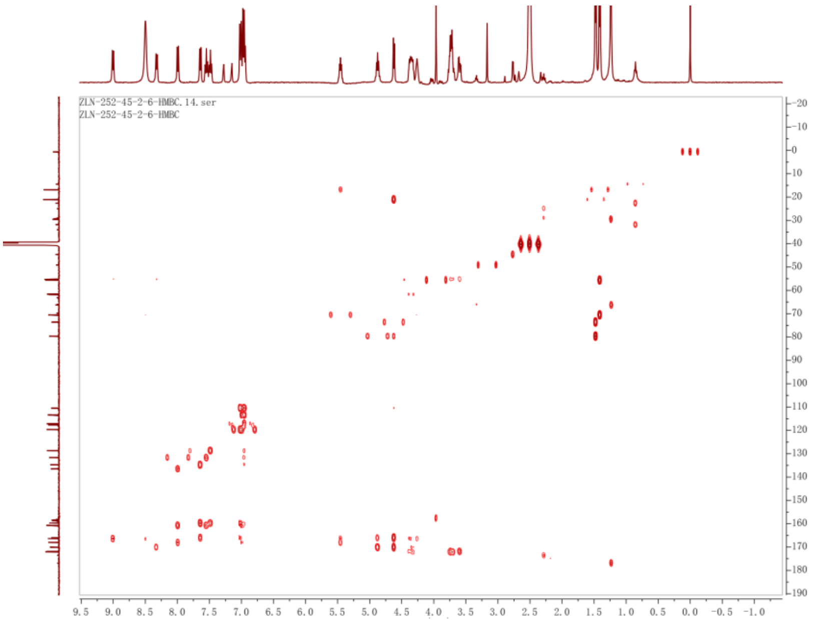


**Figure 37**. $^1$H NMR (400 MHz) spectrum of **8** in DMSO-$d_6$ ($\delta$ in ppm, $J$ in Hz)

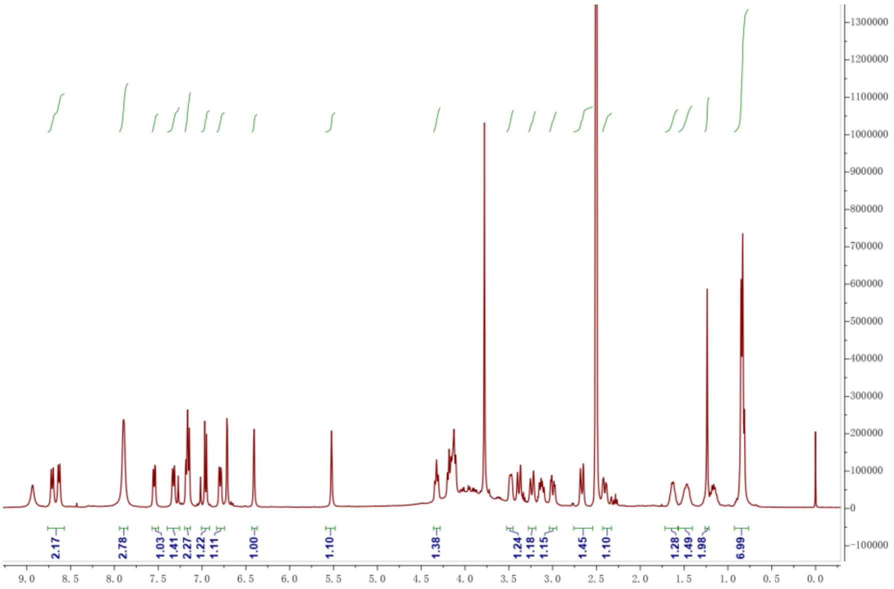


**Figure 38**. $^{13}$C NMR (125 MHz) spectrum of **8** in DMSO-$d_6$ ($\delta$ in ppm, $J$ in Hz)

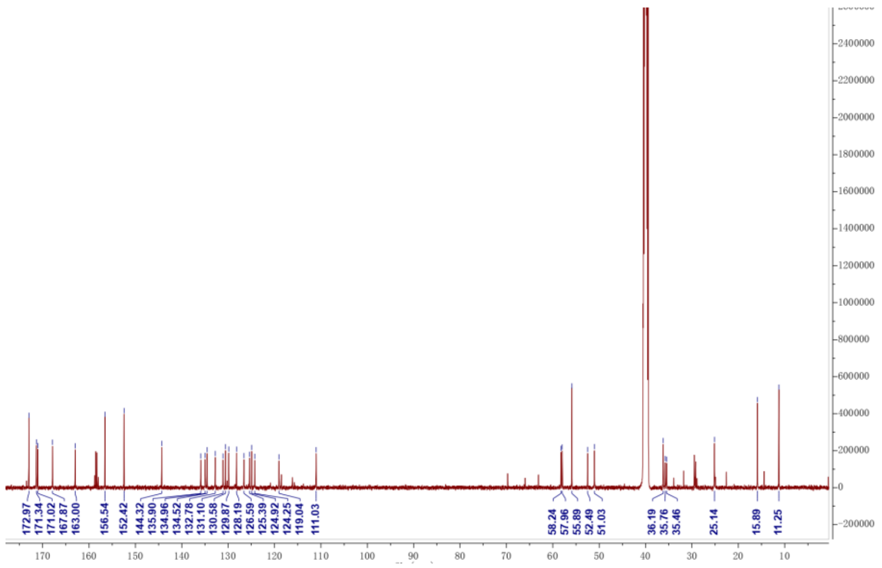


**Figure 39**. HSQC spectrum of **8** in DMSO-$d_6$ ($\delta$ in ppm, $J$ in Hz)

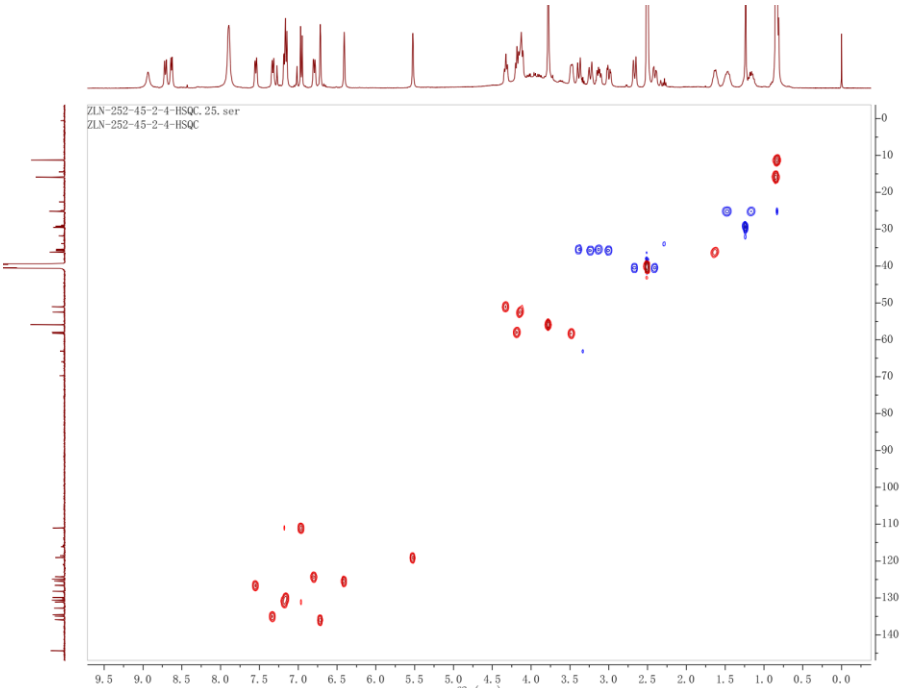


**Figure 40**. HMBC spectrum of **8** in DMSO-$d_6$ ($\delta$ in ppm, $J$ in Hz)

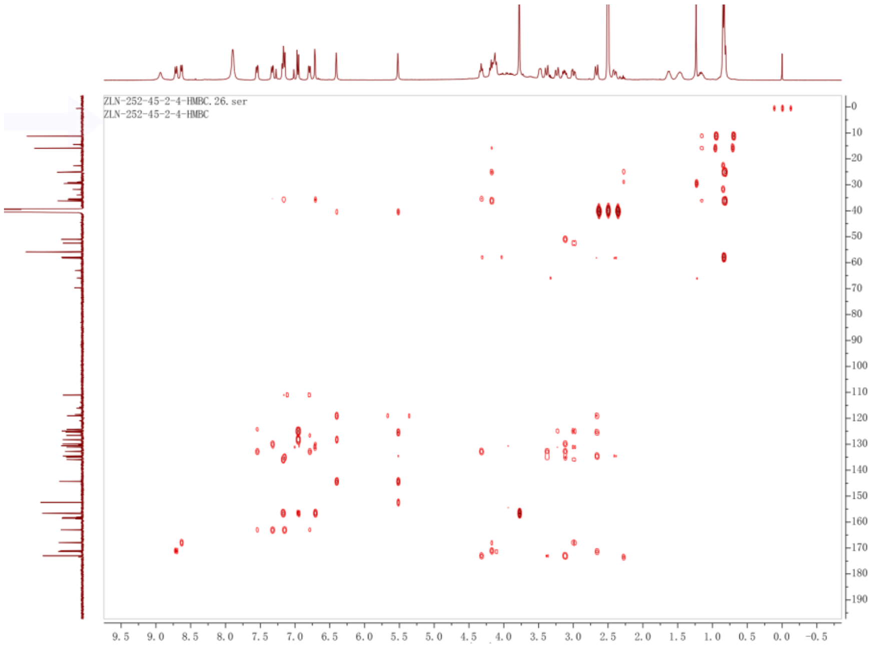


**Figure 41**. TOCSY spectrum of **8** in DMSO-$d_6$ ($\delta$ in ppm, $J$ in Hz)

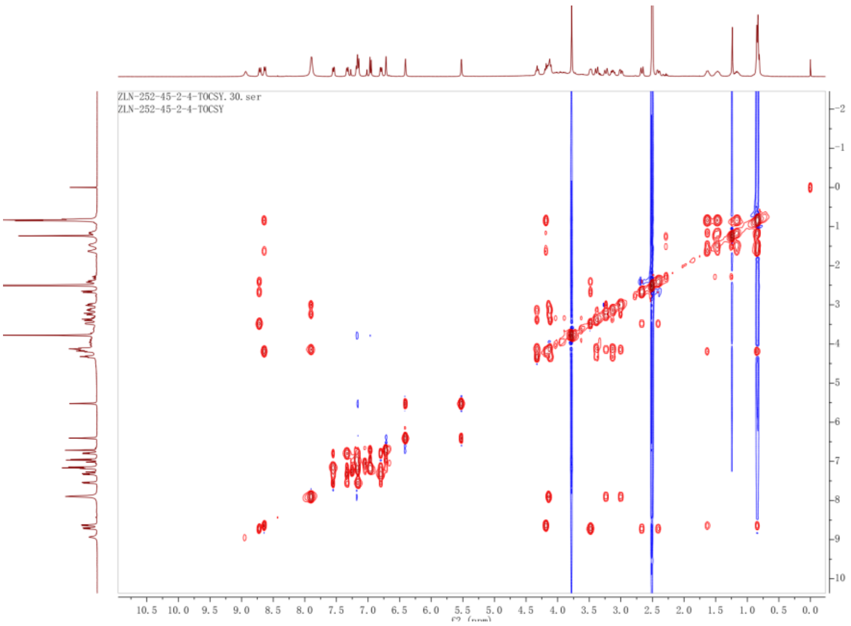


**Figure 41**. NOESY spectrum of **8** in DMSO-$d_6$ ($\delta$ in ppm, $J$ in Hz)

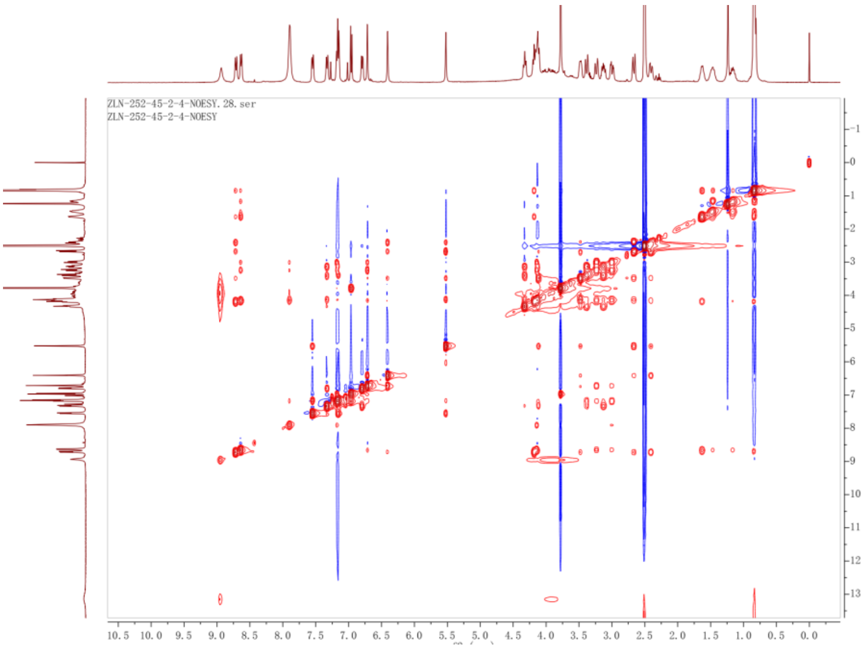
